## Supplementary material for "Profiling variable-number tandem repeat variation across populations using repeat-pangenome graphs": VNTR locus statistics (README)

chr: Chromosome where the VNTR is annotated
start: Start coordinate of the VNTR in hg38
end: End coordinate of the VNTR in hg38
ref.len: VNTR length in hg38 after boundary expansion
asm.len: Mean VNTR length across assemblies after boundary expansion
asm.len.std: Standard deviation of VNTR lengths across assemblies after boundary expansion
asm.n: Number of supporting genomes beyond hg38
hg38.acc: Genotyping accuracy using repeat-hg38 graphs measured by mean absolute
          percentage error in VNTR length. Accuracy is averaged across 16 
          unrelated genomes.
rpgg.acc: Genotyping accuracy using repeat-pangenome graphs measured by mean 
          absolute percentage error in VNTR length. Accuracy is averaged across 
          16 unrelated genomes.
qual: Averaged alignment quality of the locus measured by R-squared of the 
      linear fit between assembly and read k-mer counts. Quality score is 
      averaged across 19 genomes each using its own repeat-genome graphs.
len.vst: Vst statistic for VNTR length across the 5 populations
len.p: ANOVA P-value of all populations having a same mean in VNTR length
mikmer.r2: The variance explained in VNTR length by the most informative kmer
mikmer.ca: The most informative canonical kmer wrt VNTR length
mikmer.ca’: Reverse-complement of the most informative canonical kmer wrt VNTR 
            length
eqtl: Is an eVNTR
eqtl.n: Number of tissues with this eVNTR
eqtl.info: Association results for each discovery. 
           FORMAT: GENE1:BETA1:QVAL1,GENE2:BETA2;QVAL2[,...](TISSUE1)[;...]
