## Supplementary information for "Profiling variable-number tandem repeat variation across populations using repeat-pangenome graphs"

### Table of contents

Supplementary Figure 1. An example VNTR annotation split by adVNTR-NN. **2**

Supplementary Figure 2. Completeness of VNTR annotations in individual genomes. **3**

Supplementary Figure 3. Classes of VNTRs removed by alignment quality filtering. **5**

Supplementary Figure 4. Read sampling bias at repetitive regions. **6**

Supplementary Figure 5. Read sampling bias at non-repetitive regions of all genotyped samples. **7**

Supplementary Figure 6. Read sampling bias at non-repetitive regions preserves the relation between samples at repetitive regions. **7**

Supplementary Figure 7. Nearest neighbor search for LSB at VNTR regions using LSB at nonrepetitive regions as a proxy. **8**

Supplementary Figure 8. Profile of prediction accuracy for each sample. **9**

Supplementary Figure 9. Performance of per-locus length prediction accuracy relative to GRCh38. **10**

Supplementary Figure 10. Null and observed distributions of  and  between the EAS and AFR populations. **11**

Supplementary Figure 11. Distance of TRs and eTRs to telomere. **11**

Supplementary Figure 12. Association between the top 50 pairs of eVNTR and eGene. **12**

Supplementary Figure 13. Spurious alignment of Illumina reads to GRCh38 at a VNTR locus. **16**

Supplementary Figure 14. Boundary expansion recovers the proper boundary of TR alleles. **18**

Supplementary Figure 15. Comparison between the TR database in GangSTR and this work. **20**

Supplementary Figure 16. Distribution of number of genes overlapping shuffled high VST loci. **21**

Supplementary Figure 17. Distribution of genes and UTR regions overlapping shuffled unstable loci. **21**

Supplementary Figure 18. Number of eVNTRs shared between or specific to each tissue. **22**

Supplementary Figure 19. Length distribution of VNTRs and eVNTRs. **23**

Supplementary Figure 20. Sample QC on VNTR genotypes of the 1000 Genomes. **24**

Supplementary Figure 21. Sample QC on VNTR genotypes the GTEx Genomes. **25**

Supplementary Figure 22. Growth of relative VNTR-graph size. **26**

Supplementary Figure 23. Example of deviation in read sampling bias across samples. **26**

Supplementary Figure 24. Example of under-alignment of orthologous VNTR sequences by pggb. **27**

Supplementary Figure 25. Misalignment of simulated VNTR reads by bwa. **27**

Supplementary Figure 26. Misalignment of VNTR reads to GRCh38 rescued by danbing-tk. **28**

Supplementary Figure 27. Relationship between GC content and length prediction error. **28**

Supplementary Figure 28. Relationship between VNTR length and prediction error. **29**

Supplementary Figure 29. Relationship between eVNTR P-value and prediction error. **29**

Supplementary Methods **30**

Properties of VST 30

Supplementary Table 1. List of HGSVC members. **30**

**Supplementary Figure 1. An example VNTR annotation split by adVNTR-NN.**

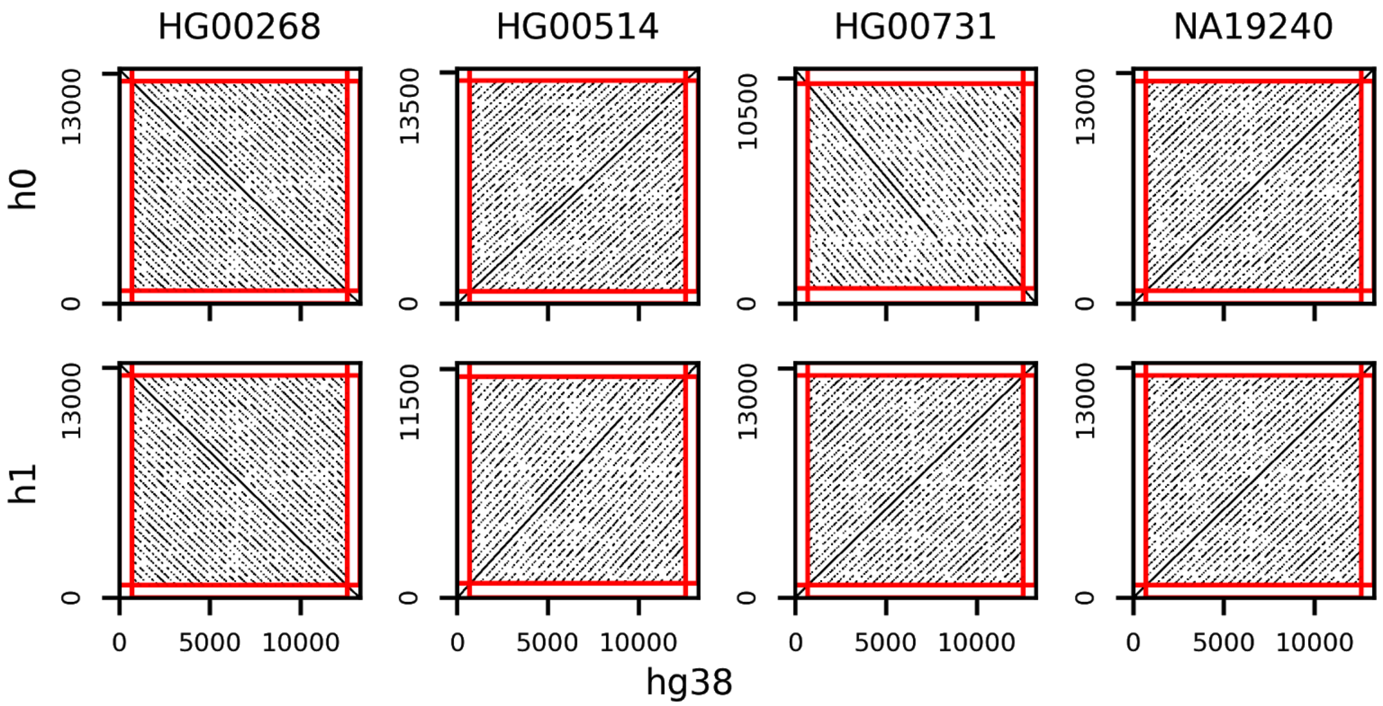


Dot plots of VNTR sequences at chr14:104941587-104953440 from four assemblies and GRCh38 are shown. Note that this region is split into 39 sub-regions in adVNTR-NN with an average VNTR size of 54 bp.

**Supplementary Figure 2. Completeness of VNTR annotations in individual genomes.**

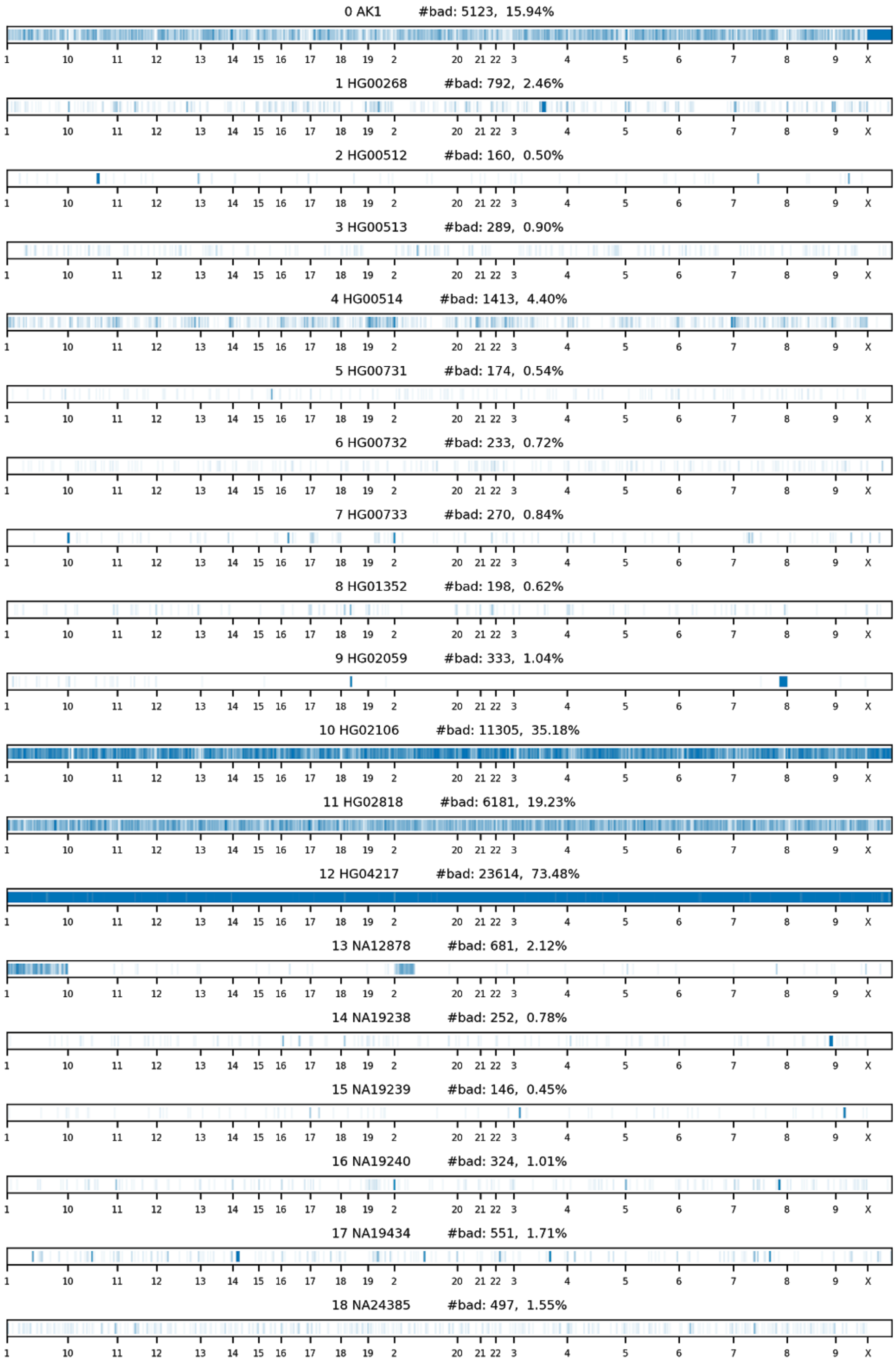

X-axis indicates relative genomic order of each missing VNTR locus which is marked by a blue stripe. A locus is called missing if both VNTR haplotypes in the genome are missing. Percentage of missing loci is the number of missing loci divided by 32,138, the total number of loci annotated.

**Supplementary Figure 3. Classes of VNTRs removed by alignment quality filtering.**

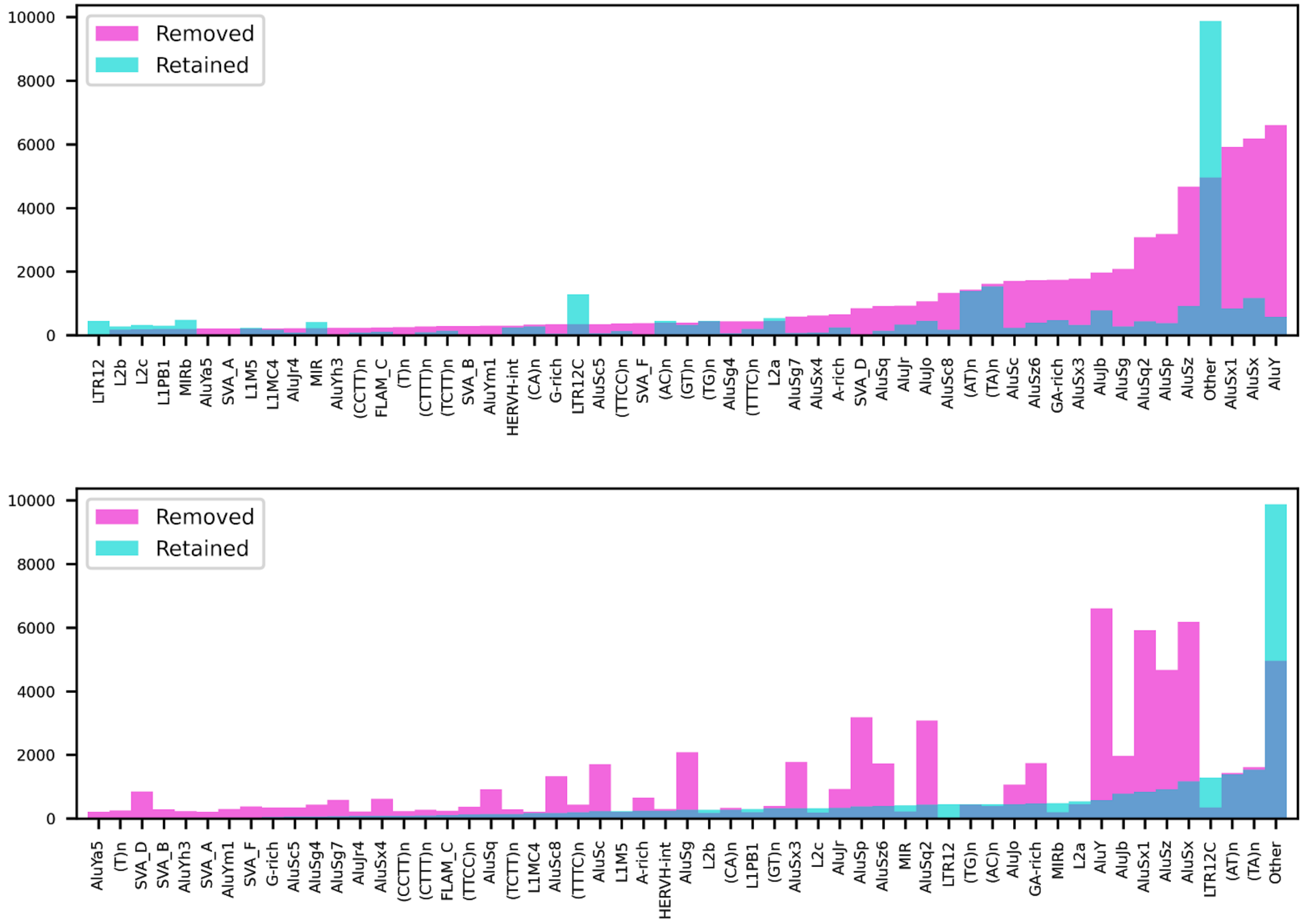


Annotations of VNTR classes are retrieved from the RepeatMasker track in UCSC Genome Browser. VNTRs that span multiple repeat annotations will be counted once for each class. Repeat classes are shown only for those with at least 200 repeats called. Class “other” indicates repeats not annotated in the RepeatMasker track. Labels on x-axis are sorted by the number of removed (top) or retained (bottom) loci.

**Supplementary Figure 4. Read sampling bias at repetitive regions.**

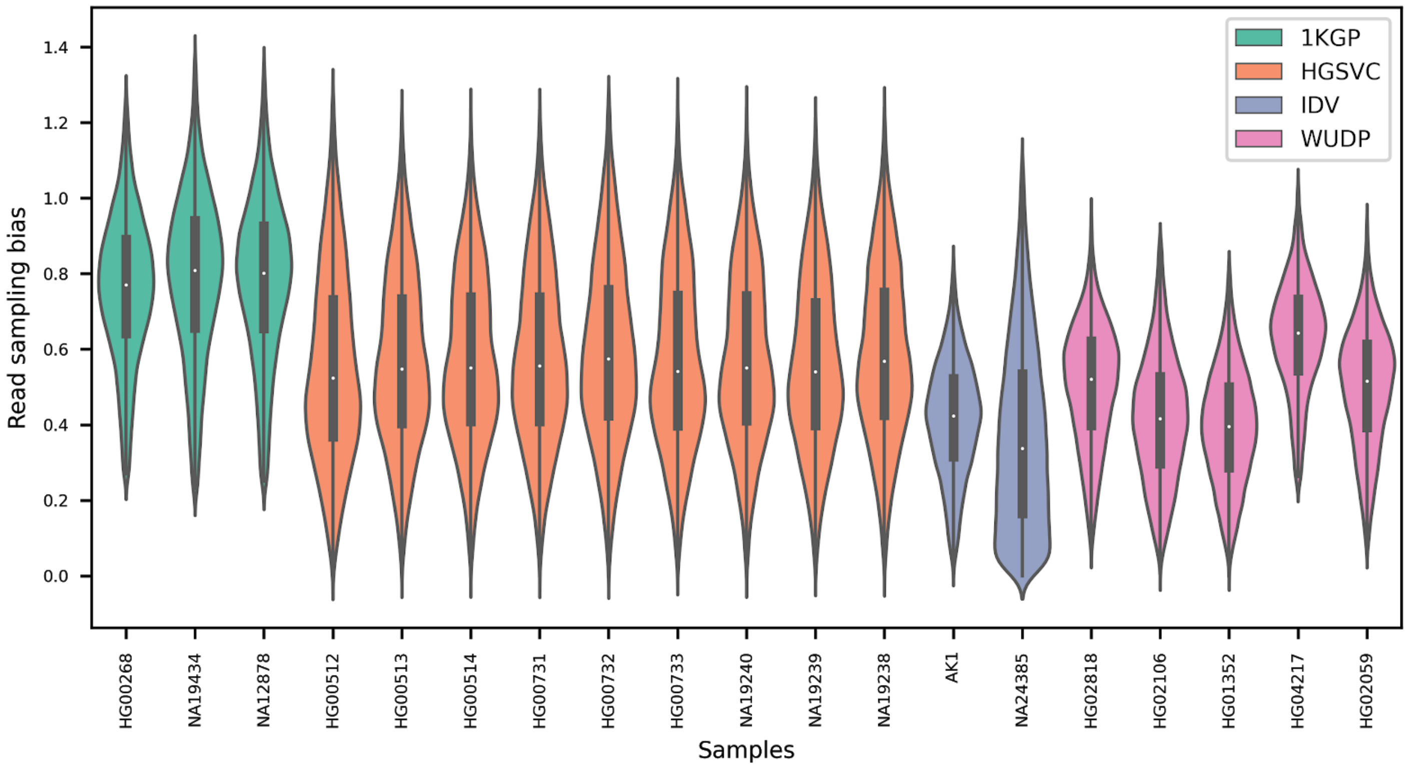


The distribution of biases at the 32,138 genotyped loci are shown for each sample. Samples are retrieved from HGSVC, Human Genome Structural Variation Consortium datasets; 1KGP, 1000 Genomes Project datasets; WUDP, Washington University Diversity Project datasets; and IDV, individual studies.

**Supplementary Figure 5. Read sampling bias at non-repetitive regions of all genotyped samples.**

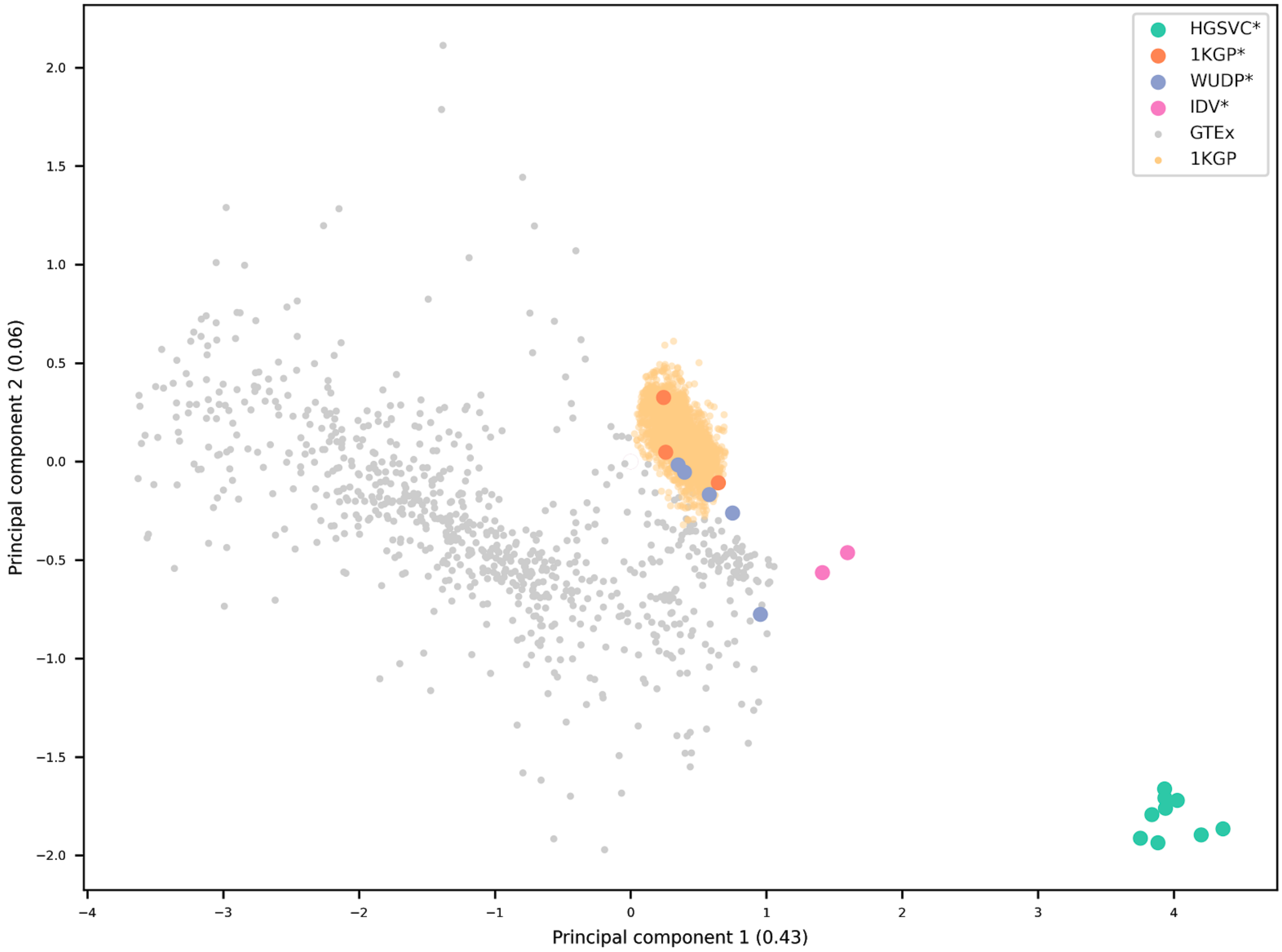


Principal component analysis was done on a $N \times L$ matrix, where $N$ is the number of samples, and $L$ is the number of unique regions. Each row of the matrix is a vector of read sampling biases in 397 unique regions from a single sample. Each sample is a tuple of (*genome*, *sequencing run*). Samples are retrieved from HGSVC, Human Genome Structural Variation Consortium datasets; 1KGP, 1000 Genomes Project datasets; WUDP, Washington University Diversity Project datasets; and IDV, individual studies; asterisks indicate samples with haplotype-resolved assemblies available.

**Supplementary Figure 6. Read sampling bias at non-repetitive regions preserves the relation between samples at repetitive regions.**

**a** **b**

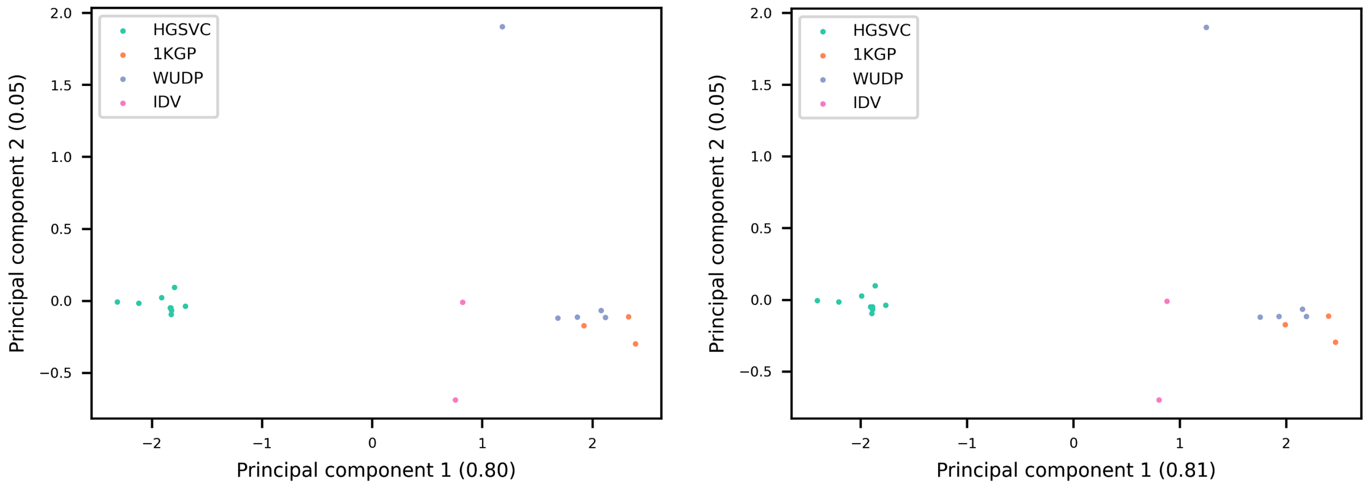


Principal component analysis of read sampling bias in VNTR (**a**) and unique (**b**) regions. PCA was done on an $N \times L$ matrix, where $N$ is the number of samples, and $L$ is the number of VNTR loci. Each row of the matrix is a vector of read sampling biases in 32,138 VNTR regions from a single sample. Each sample is a tuple of (genome, sequencing run). Samples are retrieved from HGSVC, Human Genome Structural Variation Consortium datasets; 1KGP, 1000 Genomes Project datasets; WUDP, Washington University Diversity Project datasets; and IDV, individual studies

**Supplementary Figure 7. Nearest neighbor search for LSB at VNTR regions using LSB at nonrepetitive regions as a proxy.**

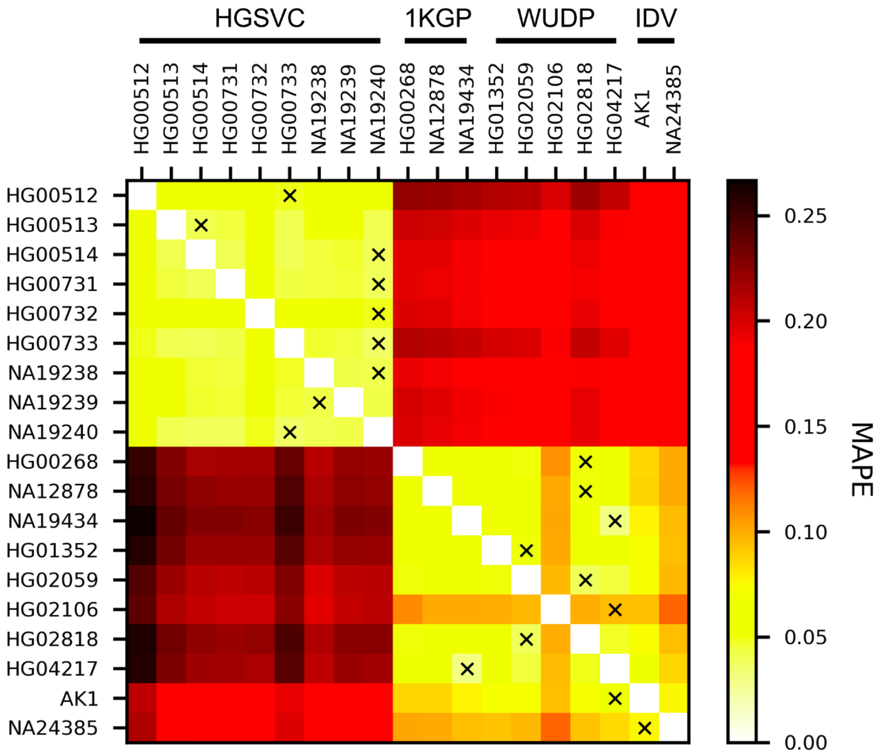

The heat map shows the mean absolute error between each pair of LSB at VNTR regions. For the sample denoted in each column, each cross indicates the nearest neighbor for that sample based on the LSB in nonrepetitive regions. HGSVC, Human Genome Structural Variation Consortium datasets; 1KGP, 1000 Genomes Project datasets; WUDP, Washington University Diversity Project datasets; IDV, individual studies.

**Supplementary Figure 8. Profile of prediction accuracy for each sample.**

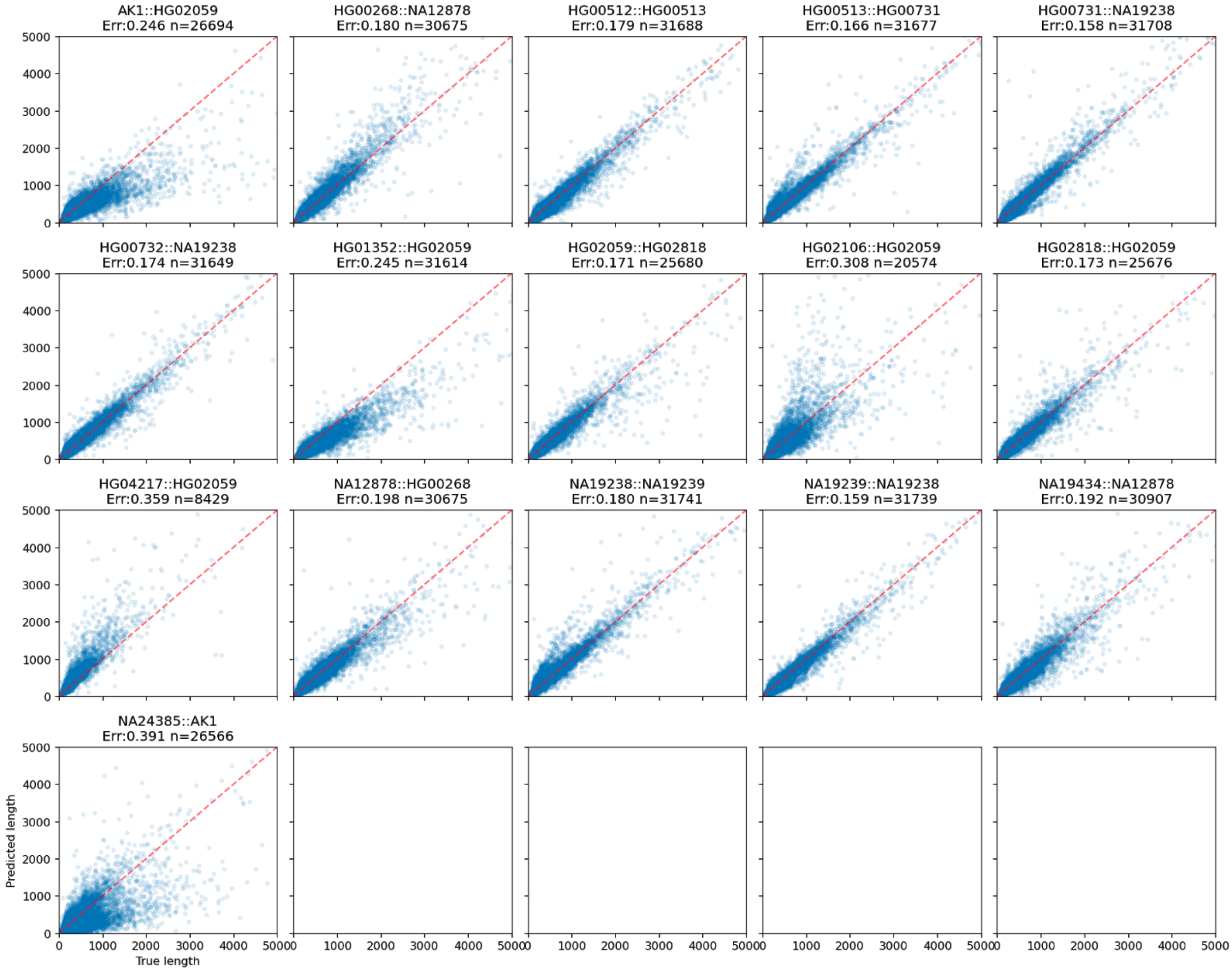


True and predicted lengths are plotted against each other for each sample. Each subtitle shows the sample name followed by its nearest sample, mean absolute percentage error and the number of loci. Loci not annotated in either the sample or its nearest sample are considered missing in the prediction step. The red dotted line shows where 100% accuracy locates.

**Supplementary Figure 9. Performance of per-locus length prediction accuracy relative to GRCh38.**

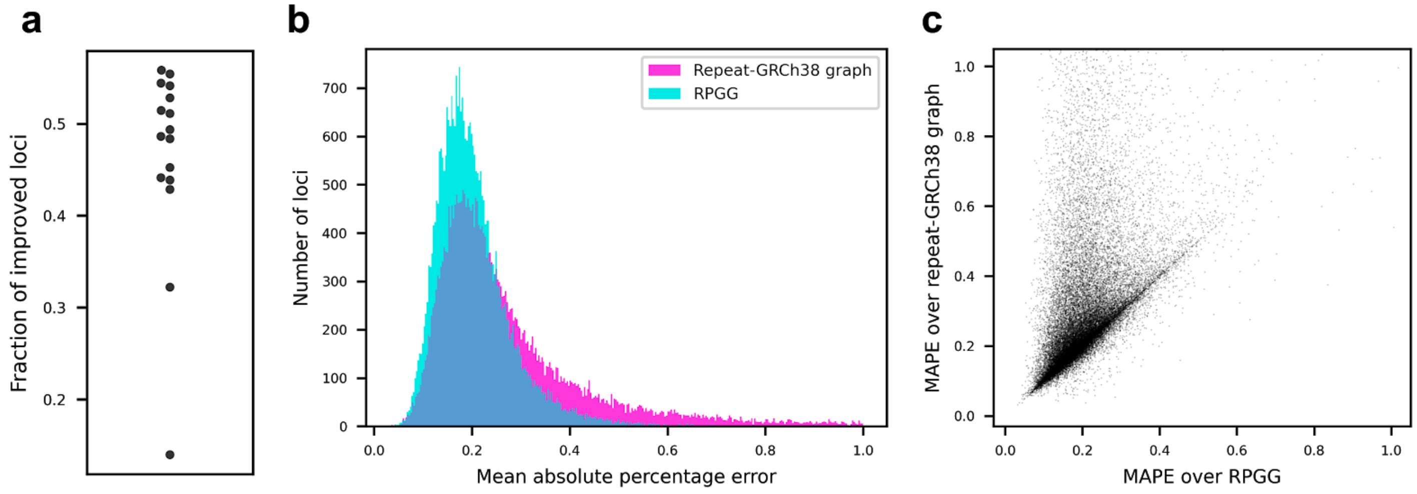


**a,** Fraction of loci with improved accuracy in each genome. **b,** Distribution of per-locus accuracy. Loci with MAPE greater than 1.0 are not shown. **c,** Per-locus MAPE of pangenome graphs versus hg38 graphs. Accuracy is measured by the mean absolute percentage error (MAPE) in VNTR lengths across all genomes (**b-c**).

**Supplementary Figure 10. Examples of unstable loci with individuals > 10 standard deviations above the mean.**

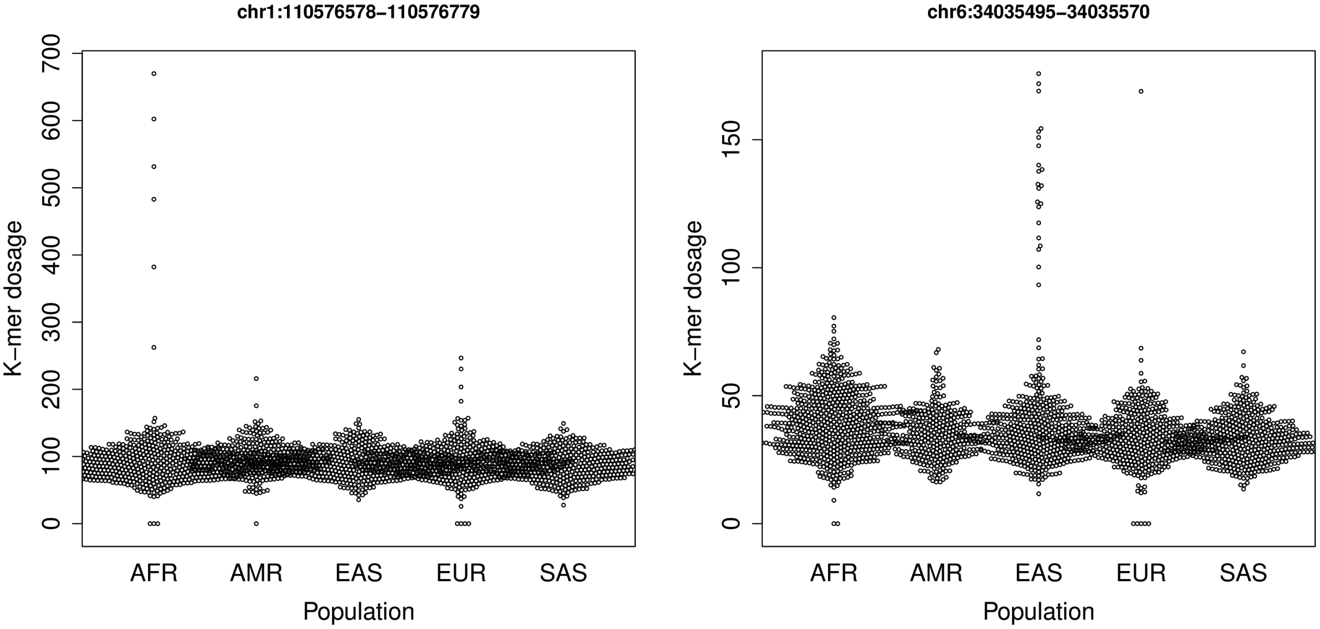


Swarm plots demonstrating highly unstable loci, determined as having an individual with coverage at least ten standard deviations above the mean. The locus on the left overlaps *KCNA2*, and the locus on the right overlaps GRM4.

**Supplementary Figure 11. Null and observed distributions of $kmc_d$ and $r_d^2$ between the EAS and AFR populations.**

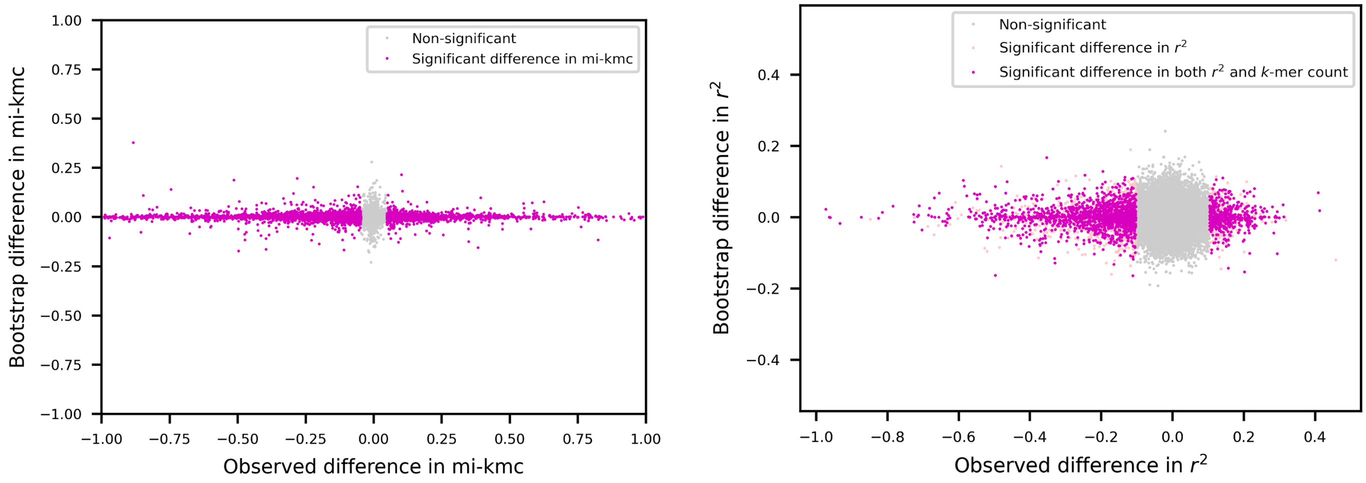


The Null distribution of difference in the count of the most informative *k*-mer (mi-kmc, left) and difference in variance explained by the most informative *k*-mer ($r^2$, right) at each locus was simulated using bootstrap from the EAS population with sample size matching the sum of both samples ($N_{EAS}$=502, $N_{AFR}$=661). Observed values within the two-tailed P<0.01 regions were called significant.

**Supplementary Figure 12. Distance of TRs and eTRs to telomere.**

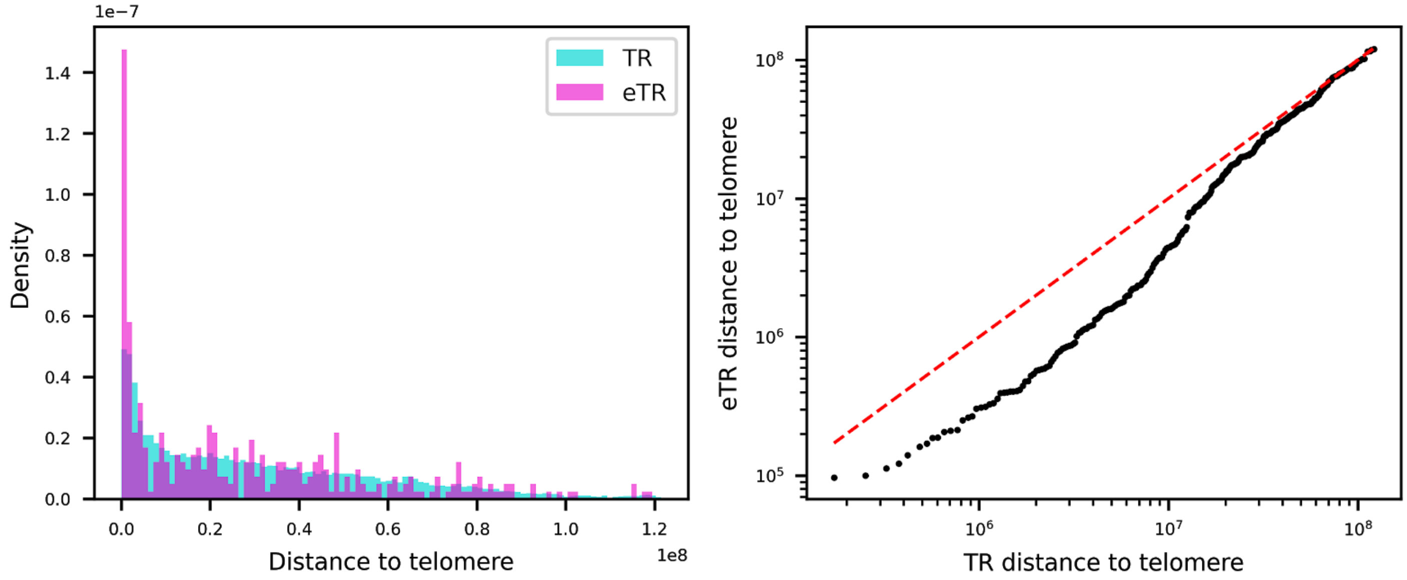


Telomere annotations were retrieved from UCSC Genome Browser and used to find the distance of a tandem repeats to its closest telomere. Distribution (left) and the q-q plot (right) of the statistics from TRs and eTRs were compared.

**Supplementary Figure 13. Association between the top 50 pairs of eVNTR and eGene.**

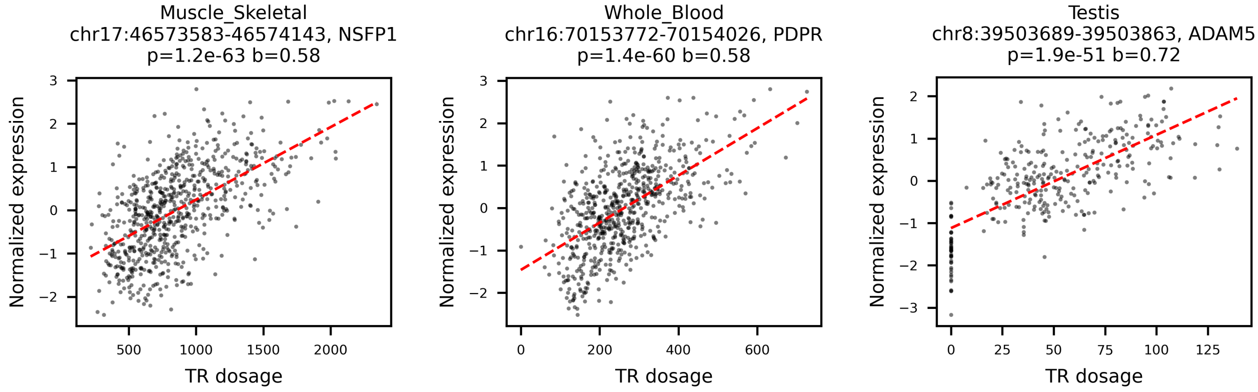

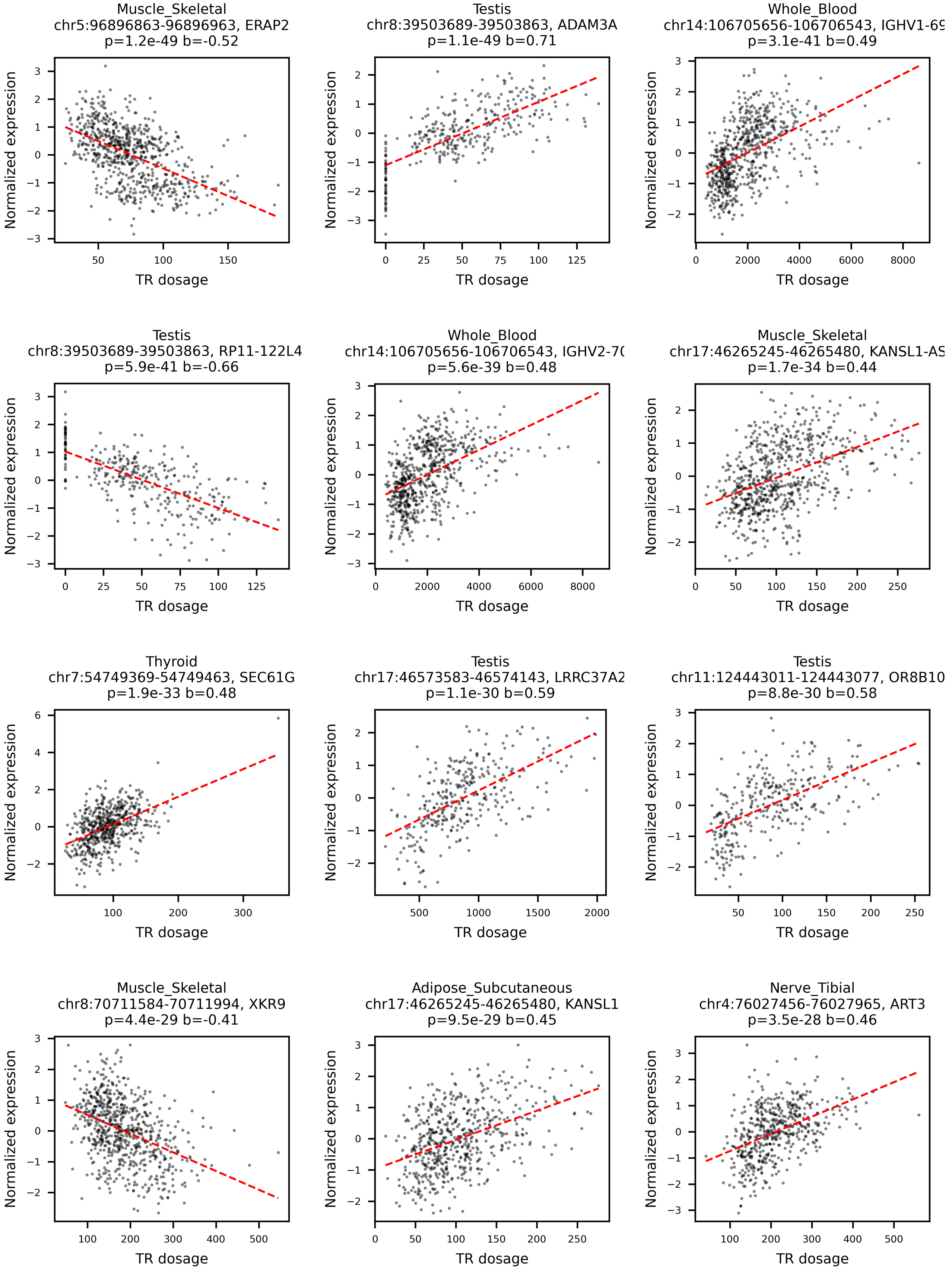
Muscle_Skeletal
chr5:96896863-96896963, ERAP2
p=1.2e-49 b=-0.52
Testis
chr8:39503689-39503863, ADAM3A
p=1.1e-49 b=0.71
Whole_Blood
chr14:106705656-106706543, IGHV1-69
p=3.1e-41 b=0.49
Testis
chr8:39503689-39503863, RP11-122L4
p=5.9e-41 b=-0.66
Whole_Blood
chr14:106705656-106706543, IGHV2-70
p=5.6e-39 b=0.48
Muscle_Skeletal
chr17:46265245-46265480, KANSL1-AS
p=1.7e-34 b=0.44
Thyroid
chr7:54749369-54749463, SEC61G
p=1.9e-33 b=0.48
Testis
chr17:46573583-46574143, LRRC37A2
p=1.1e-30 b=0.59
Testis
chr11:124443011-124443077, OR8B10
p=8.8e-30 b=0.58
Muscle_Skeletal
chr8:70711584-70711994, XKR9
p=4.4e-29 b=-0.41
Adipose_Subcutaneous
chr17:46265245-46265480, KANSL1
p=9.5e-29 b=0.45
Nerve_Tibial
chr4:76027456-76027965, ART3
p=3.5e-28 b=0.46
Normalized expression
TR dosage

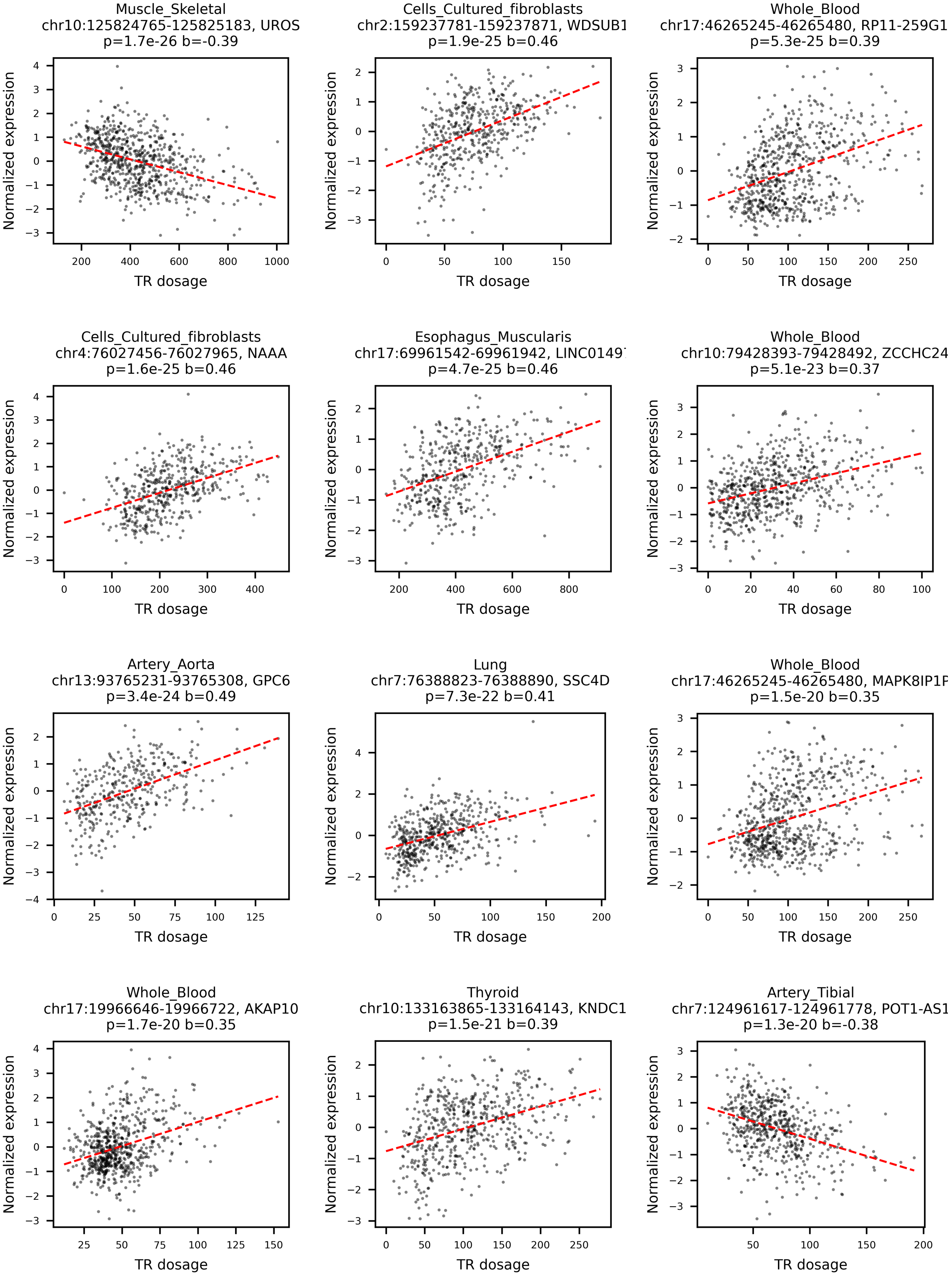
Muscle_Skeletal
chr10:125824765-125825183, UROS
p=1.7e-26 b=-0.39
Cells_Cultured_fibroblasts
chr2:159237781-159237871, WDSUB1
p=1.9e-25 b=0.46
Whole_Blood
chr17:46265245-46265480, RP11-259G1
p=5.3e-25 b=0.39
Cells_Cultured_fibroblasts
chr4:76027456-76027965, NAAA
p=1.6e-25 b=0.46
Esophagus_Muscularis
chr17:69961542-69961942, LINC0149
p=4.7e-25 b=0.46
Whole_Blood
chr10:79428393-79428492, ZCCHC24
p=5.1e-23 b=0.37
Artery_Aorta
chr13:93765231-93765308, GPC6
p=3.4e-24 b=0.49
Lung
chr7:76388823-76388890, SSC4D
p=7.3e-22 b=0.41
Whole_Blood
chr17:46265245-46265480, MAPK8IP1F
p=1.5e-20 b=0.35
Whole_Blood
chr17:19966646-19966722, AKAP10
p=1.7e-20 b=0.35
Thyroid
chr10:133163865-133164143, KNDC1
p=1.5e-21 b=0.39
Artery_Tibial
chr7:124961617-124961778, POT1-AS1
p=1.3e-20 b=-0.38
Normalized expression
TR dosage

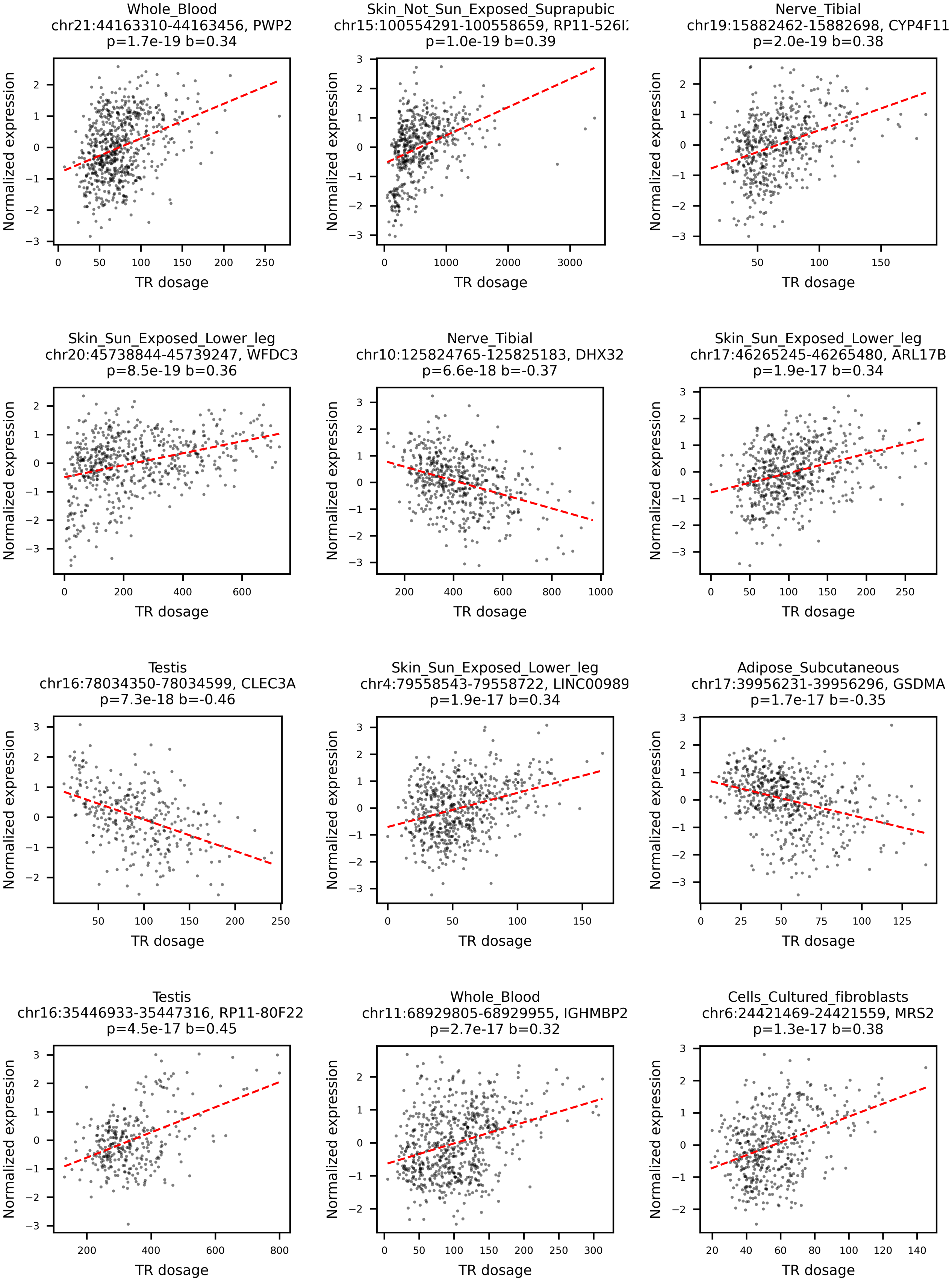
Whole_Blood
chr21:44163310-44163456, PWP2
p=1.7e-19 b=0.34
Skin_Not_Sun_Exposed_Suprapubic
chr15:100554291-100558659, RP11-526I
p=1.0e-19 b=0.39
Nerve_Tibial
chr19:15882462-15882698, CYP4F11
p=2.0e-19 b=0.38
Skin_Sun_Exposed_Lower_leg
chr20:45738844-45739247, WFDC3
p=8.5e-19 b=0.36
Nerve_Tibial
chr10:125824765-125825183, DHX32
p=6.6e-18 b=-0.37
Skin_Sun_Exposed_Lower_leg
chr17:46265245-46265480, ARL17B
p=1.9e-17 b=0.34
Testis
chr16:78034350-78034599, CLEC3A
p=7.3e-18 b=-0.46
Skin_Sun_Exposed_Lower_leg
chr4:79558543-79558722, LINC00989
p=1.9e-17 b=0.34
Adipose_Subcutaneous
chr17:39956231-39956296, GSDMA
p=1.7e-17 b=-0.35
Testis
chr16:35446933-35447316, RP11-80F22
p=4.5e-17 b=0.45
Whole_Blood
chr11:68929805-68929955, IGHMBP2
p=2.7e-17 b=0.32
Cells_Cultured_fibroblasts
chr6:24421469-24421559, MRS2
p=1.3e-17 b=0.38
Normalized expression
TR dosage

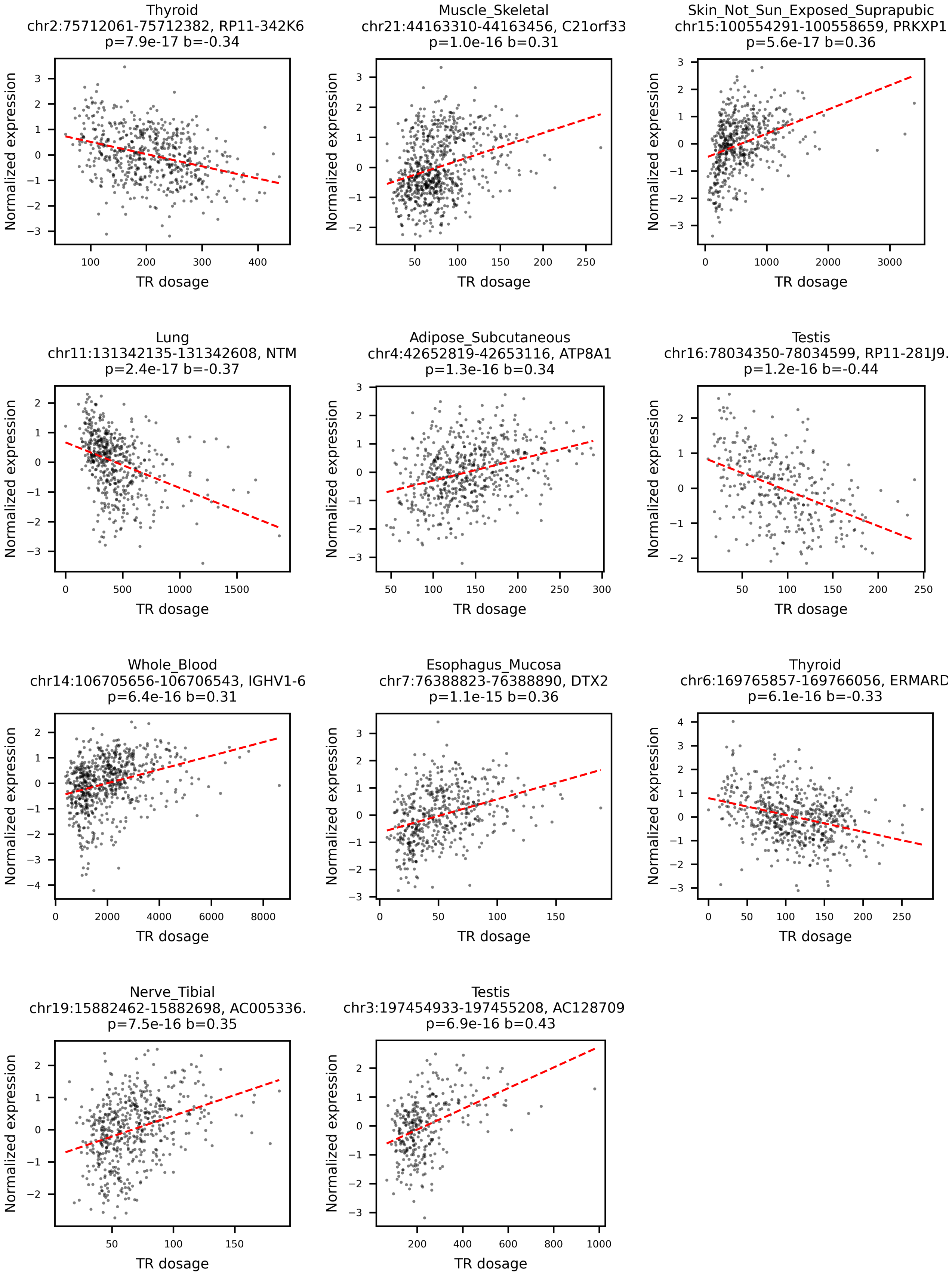
Thyroid
chr2:75712061-75712382, RP11-342K6
p=7.9e-17 b=-0.34
Muscle_Skeletal
chr21:44163310-44163456, C21orf33
p=1.0e-16 b=0.31
Skin_Not_Sun_Exposed_Suprapubic
chr15:100554291-100558659, PRKXP1
p=5.6e-17 b=0.36
Lung
chr11:131342135-131342608, NTM
p=2.4e-17 b=-0.37
Adipose_Subcutaneous
chr4:42652819-42653116, ATP8A1
p=1.3e-16 b=0.34
Testis
chr16:78034350-78034599, RP11-281J9.
p=1.2e-16 b=-0.44
Whole_Blood
chr14:106705656-106706543, IGHV1-6
p=6.4e-16 b=0.31
Esophagus_Mucosa
chr7:76388823-76388890, DTX2
p=1.1e-15 b=0.36
Thyroid
chr6:169765857-169766056, ERMARD
p=6.1e-16 b=-0.33
Nerve_Tibial
chr19:15882462-15882698, AC005336.
p=7.5e-16 b=0.35
Testis
chr3:197454933-197455208, AC128709
p=6.9e-16 b=0.43
Normalized expression
TR dosage

Plots are shown in order of q-value. The format of plot titles is tissue, VNTR_region, gene_name, nominal_p_val and effect_size. The linear fit is shown as a dashed red line.

**Supplementary Figure 14. Spurious alignment of Illumina reads to GRCh38 at a VNTR locus.**

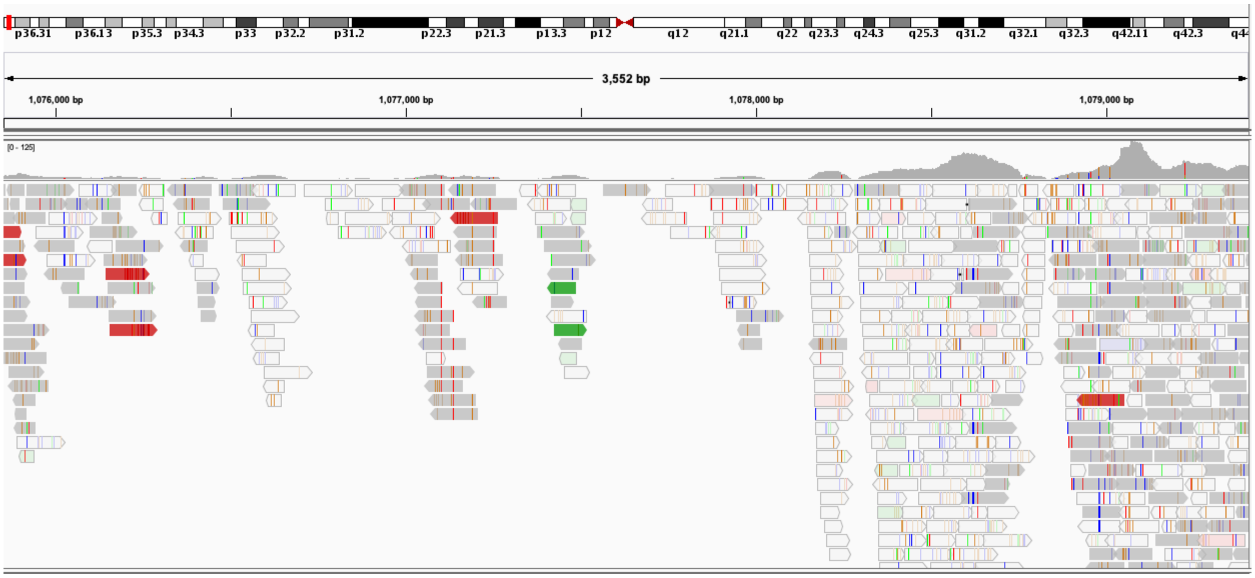


Alignment of Illumina datasets at 60x coverage from the HG00514 individual to chr1:1075852-1079425 of hg38 is visualized by Integrative Genomics Viewer (IGV).

**Supplementary Figure 15. Boundary expansion recovers the proper boundary of TR alleles.**

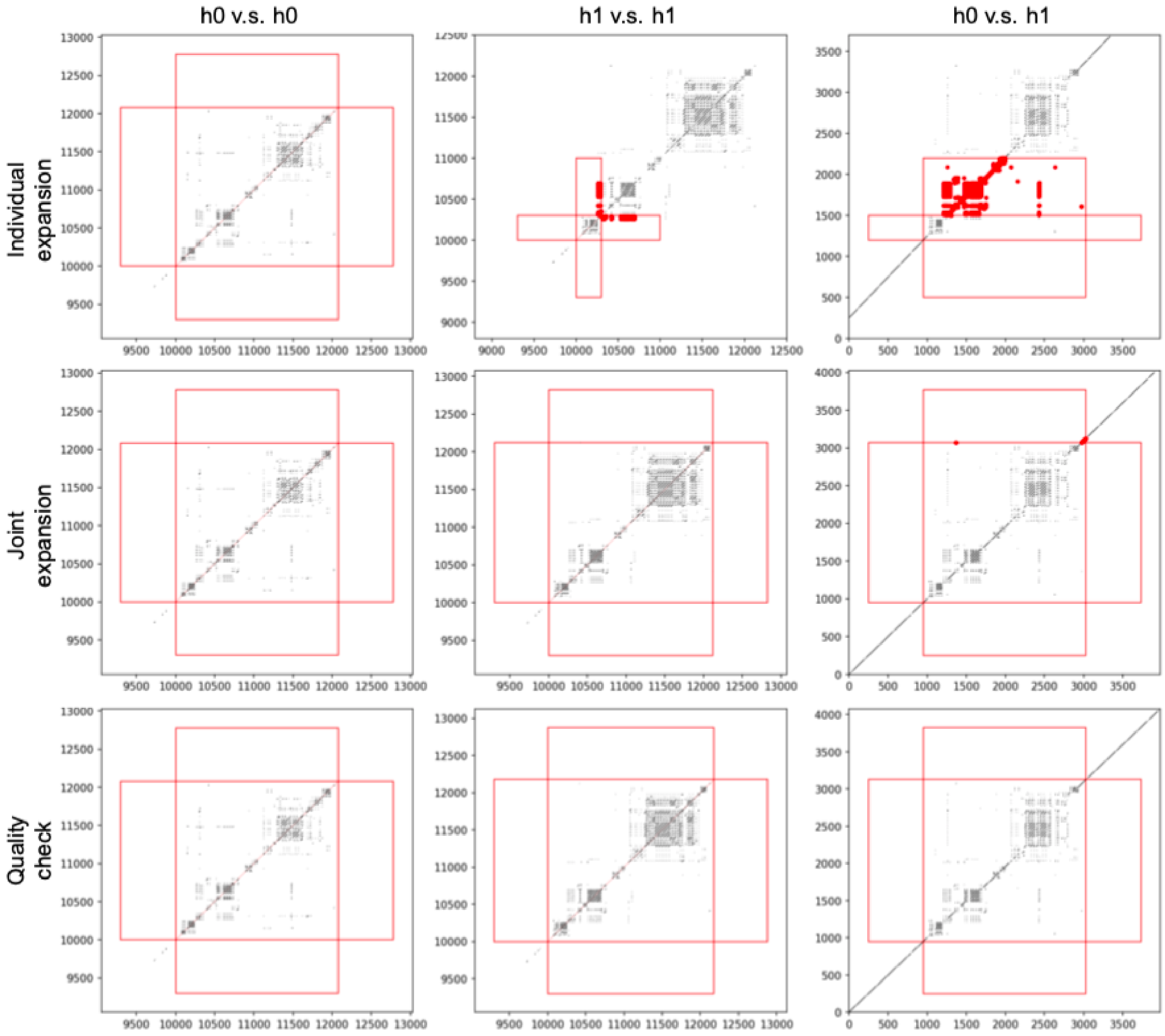


For every two TR alleles, the boundary expansion algorithm operates in three steps: individual expansion, joint expansion and quality check (Methods). The red boxes indicate the regions where *k*-mer matching is subject to inspection. Any matches (red dots) occurring outside of the central red box indicate the presence of shared *k*-mers between the TR and the flanking sequence.

**Supplementary Figure 16. Comparison between the TR database in GangSTR and this work.**

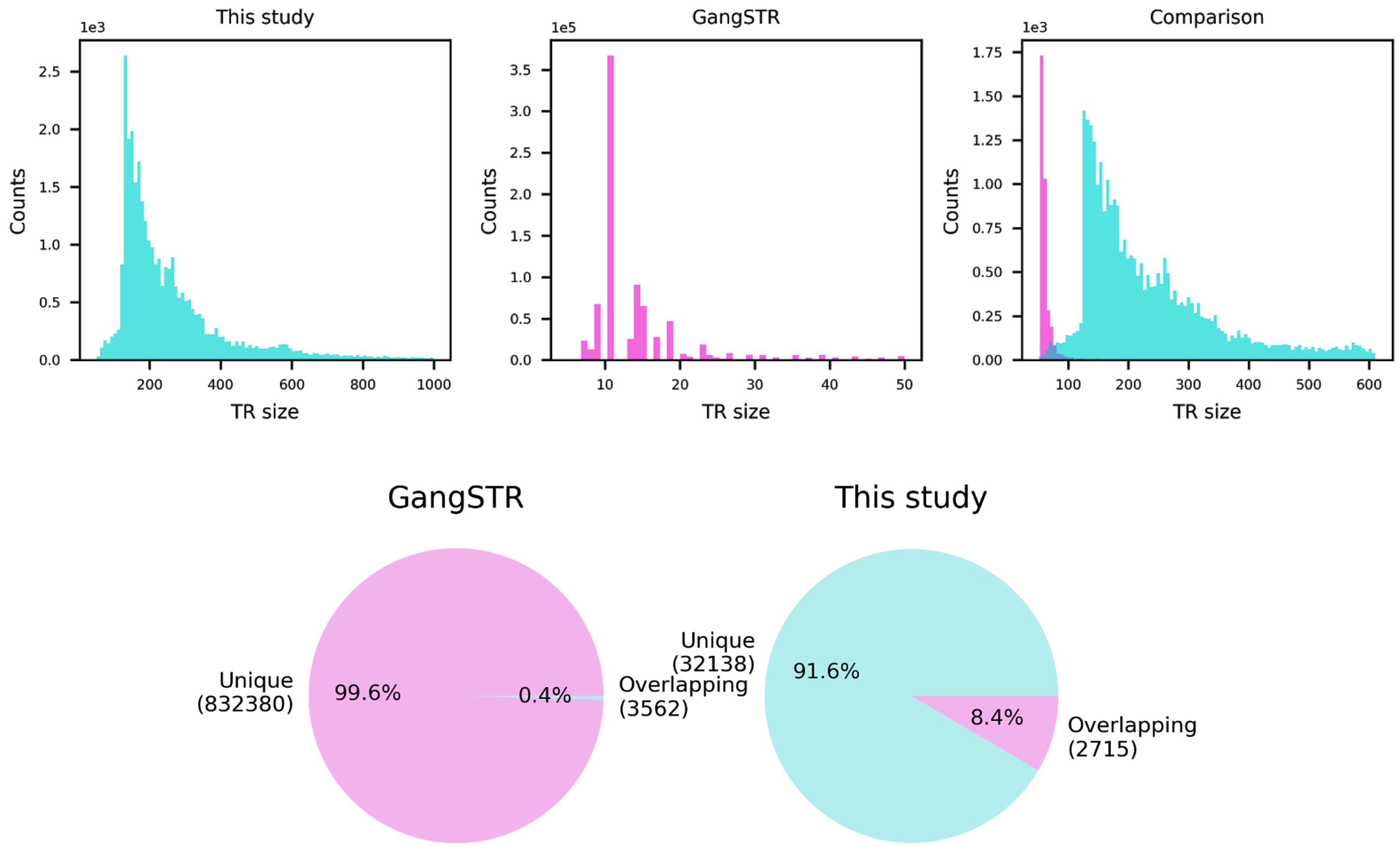


**a,** Size distribution of the TRs annotated in each study. TRs with size greater than 150 bp in at least one assembly and with size greater than 50 bp in hg38 are annotated in this study. TR sizes above 1000 bp, above 50 bp and below 50 bp are not shown for this study (left), GangSTR (middle) and comparison (right), respectively. **b,** Percentage of overlapping TRs between databases. The number of overlapping loci changes across databases since multiple loci in GangSTR's database could correspond to only one locus in our database.

**Supplementary Figure 17. Distribution of number of genes overlapping shuffled high $V_{ST}$ loci.**

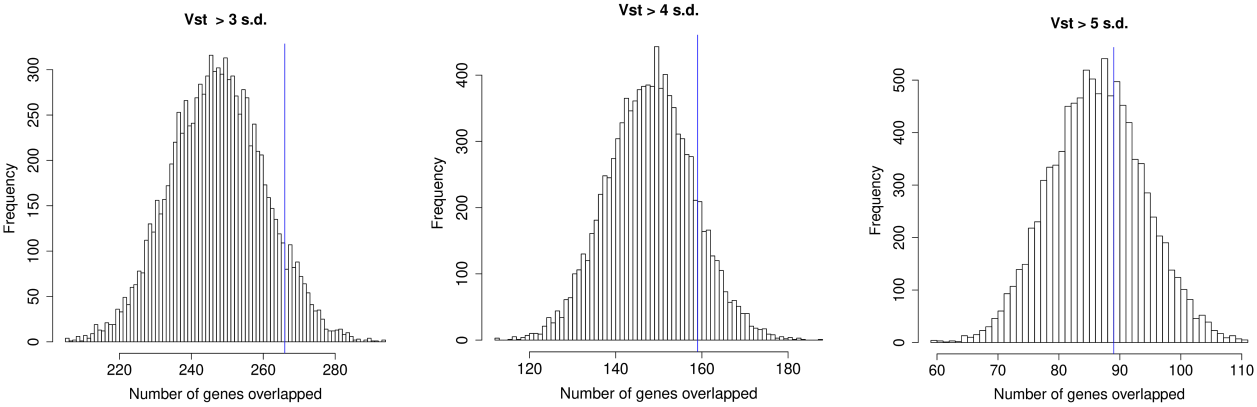


The frequency for 10,000 iterations of the number of genes overlapping high $V_{ST}$ loci that are shuffled across the euchromatic genome. High $V_{ST}$ are defined by a minimal number of standard deviations above the mean (3-5) (N=785, 470, and 235). The number of genes overlapping high $V_{ST}$ loci in the original dataset are shown by the full-height vertical lines.

**Supplementary Figure 18. Distribution of genes and UTR regions overlapping shuffled unstable loci.**

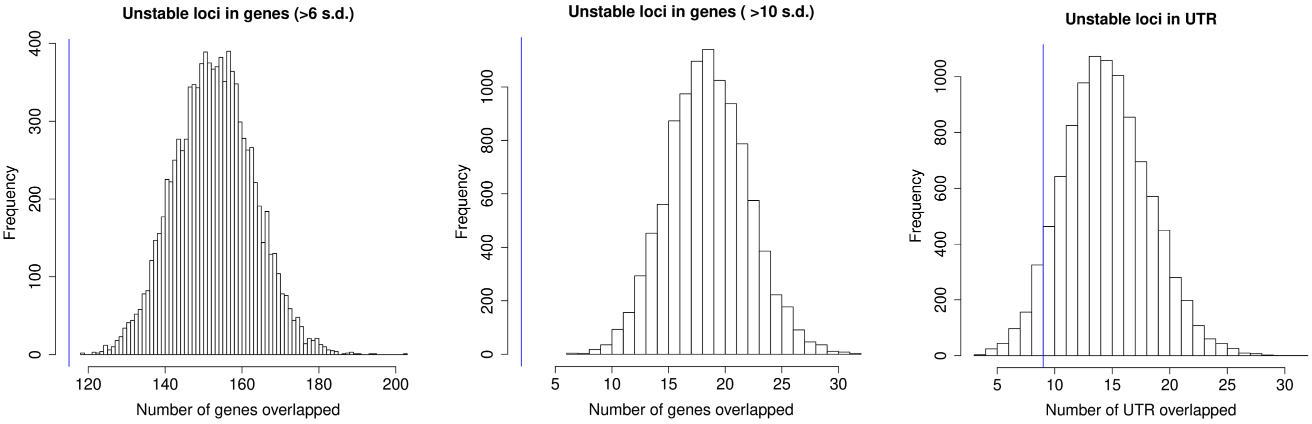


The number of genes overlapping VNTRs defined as unstable with different cutoff values: at least one individual with dosage > 6 standard deviations above the mean (N=19), and with > 10 standard deviations above the mean (N=2). The number of genes/UTRs overlapping unstable loci in the original dataset are shown by the full-height vertical lines.

**Supplementary Figure 19. Number of eVNTRs shared between or specific to each tissue.**

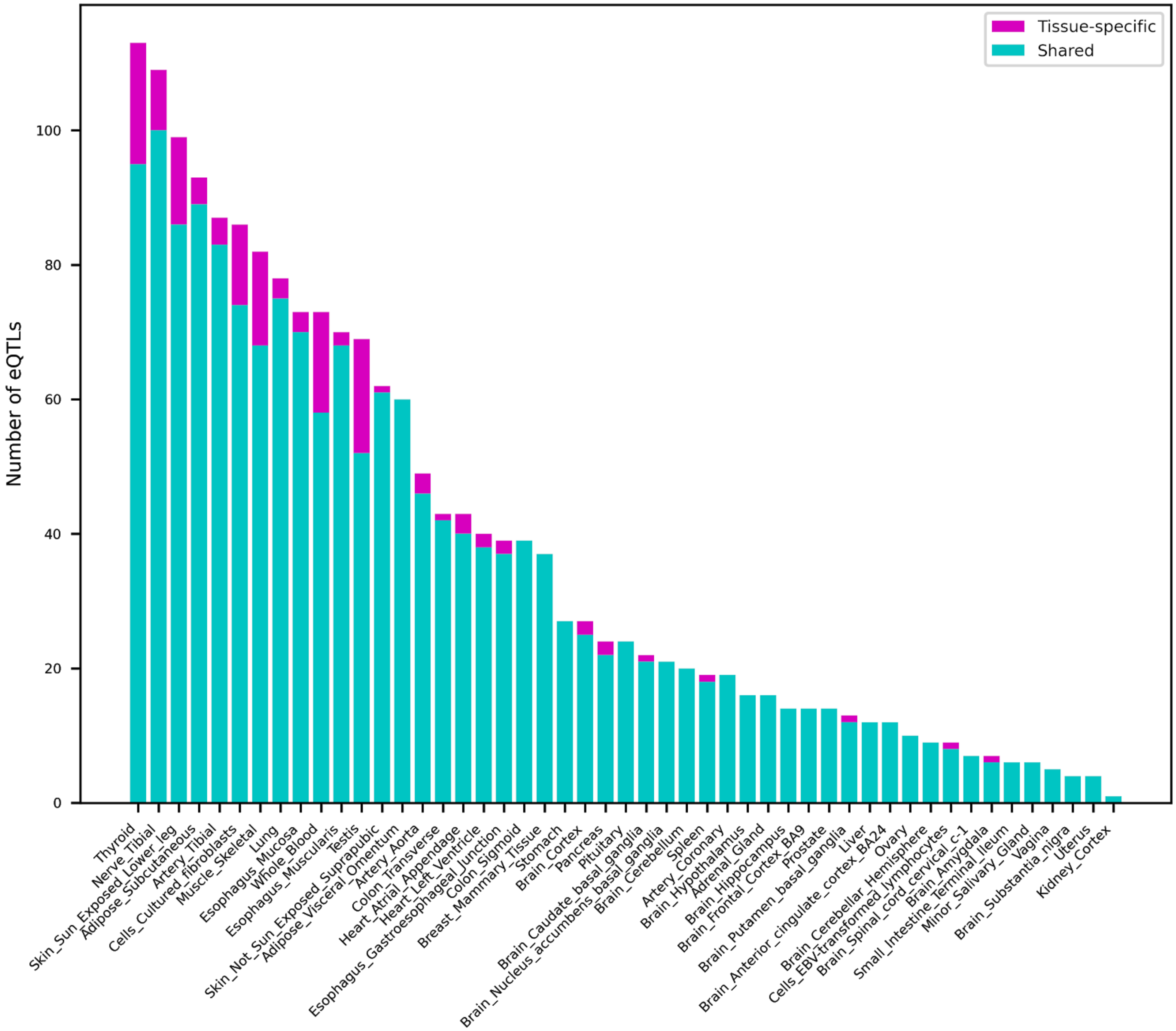


eQTL discoveries for the 32,138 VNTR loci were controlled at 5% FDR.

**Supplementary Figure 20. Length distribution of VNTRs and eVNTRs.**

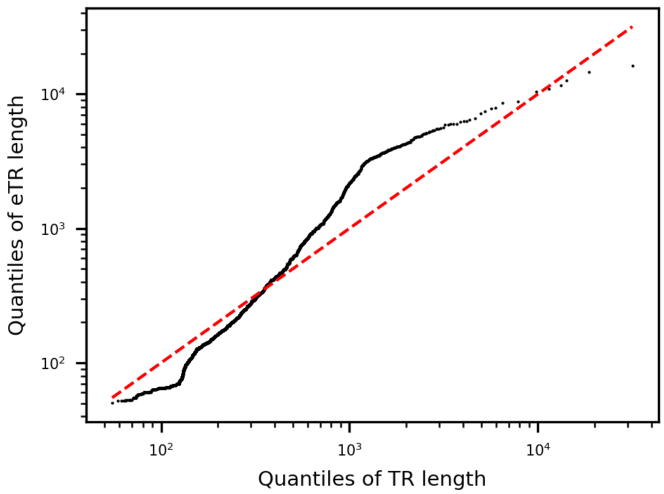


Length distribution of eVNTRs and VNTRs. eQTL discoveries for the 32,138 VNTR loci were controlled at 5% FDR.

**Supplementary Figure 21. Sample QC on VNTR genotypes of the 1000 Genomes.**

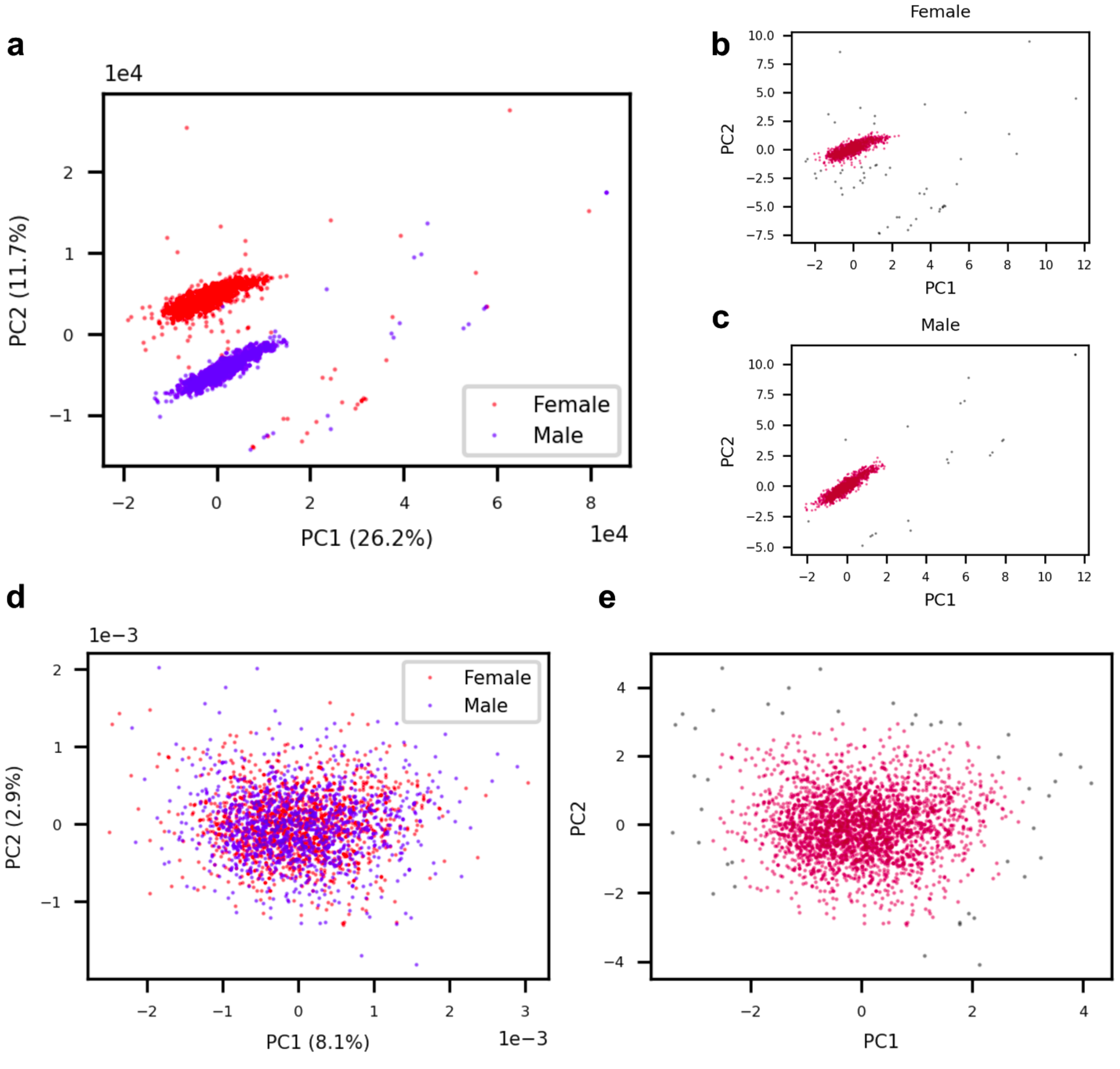


**a,** Joint PCA plot of samples using the *k*-mer dosage adjusted by coverage. **b-c,** Outlier detection, shown in gray, using DBSCAN with eps=0.5 on male (b) and female individuals (c). **d-e** Joint PCA plot of samples using the read sampling biases from 397 control regions (c) and the outliers detected using DBSCAN with eps=0.5 (d).

**Supplementary Figure 22. Sample QC on VNTR genotypes the GTEx Genomes.**

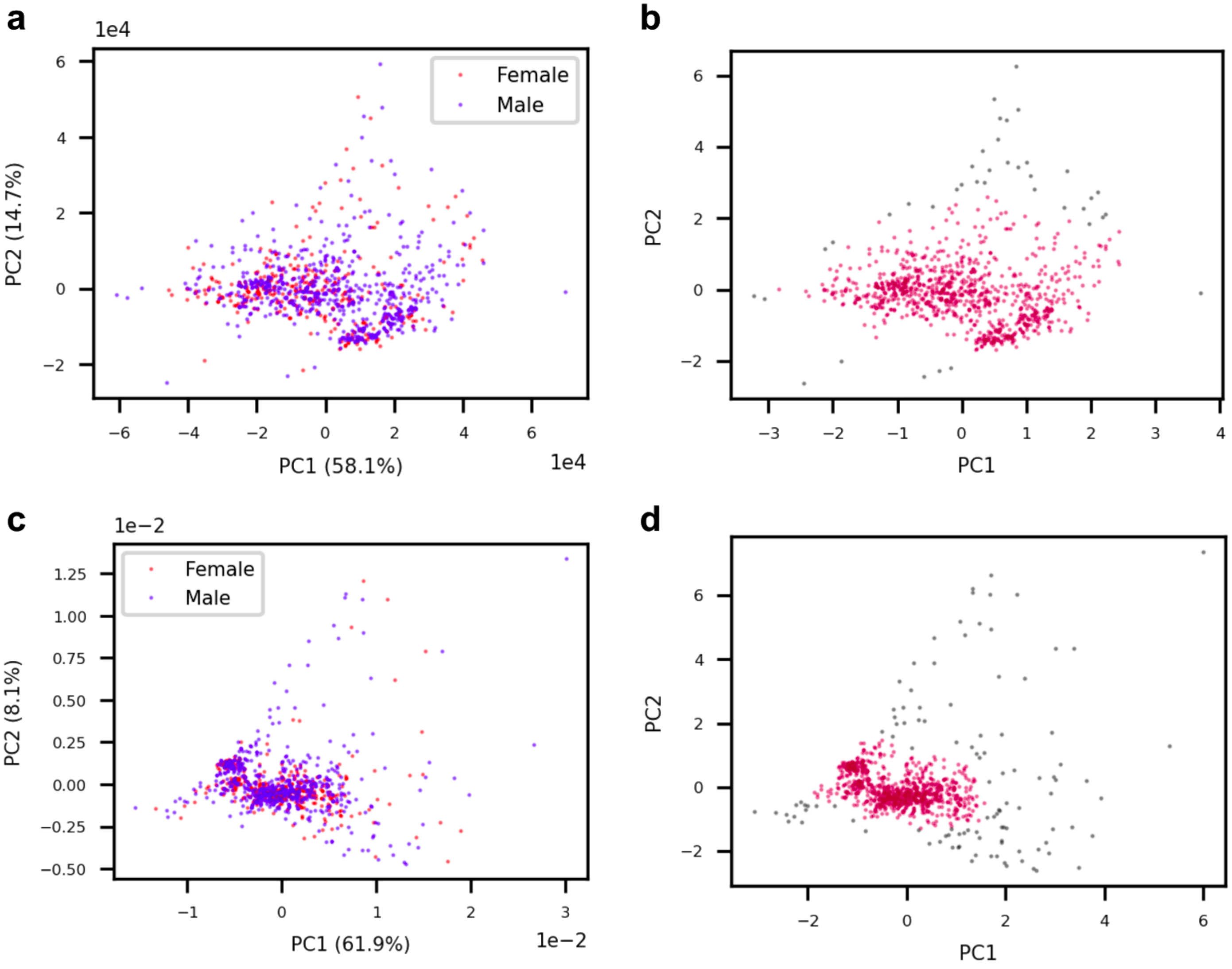


**a-b** Joint PCA plot of samples using the *k*-mer dosage adjusted by coverage and allelic dosage. (a) and the outliers detected, shown in gray, using DBSCAN with eps=0.5 (b). **c-d** Joint PCA plot of samples using the read sampling biases from 397 control regions (c) and the outliers detected using DBSCAN with eps=0.3 (d).

**Supplementary Figure 23. Growth of relative VNTR-graph size.**

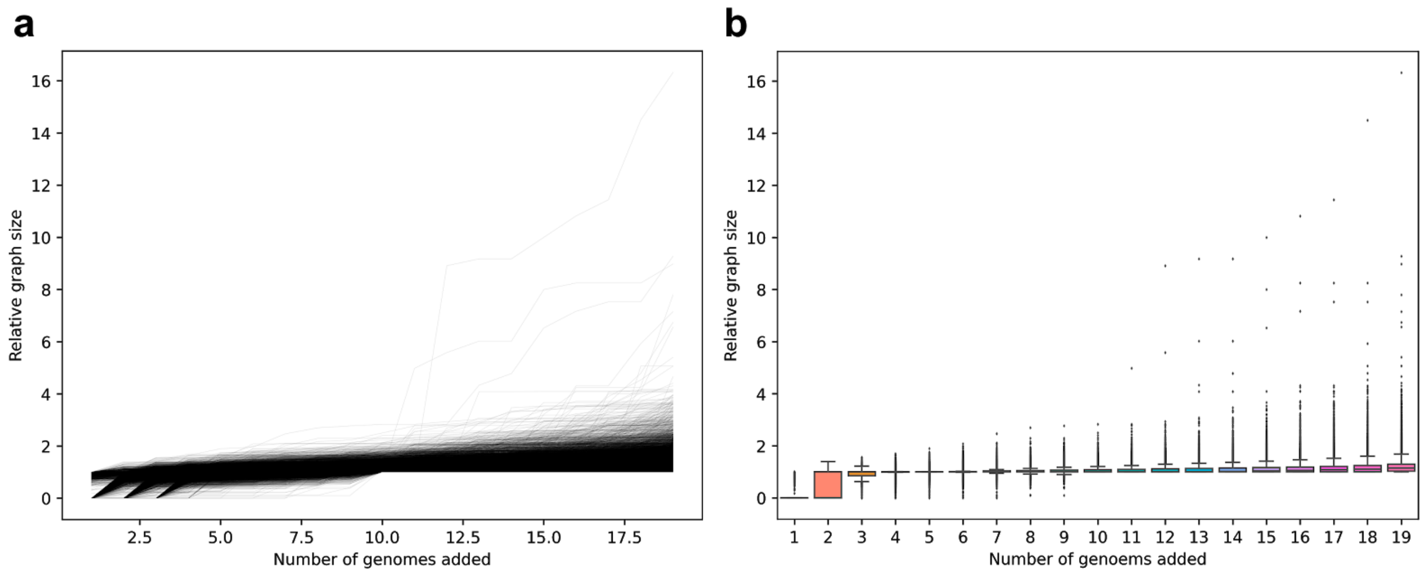


The growth curve (**a**) and the distribution of graph size (**b**) if adding genomes in an incremental manner are shown for the 32,138 VNTR loci. Relative graph size is the ratio between the number of nodes, or *k*-mers, in the RPGG and the median number of nodes in a single genome.

**Supplementary Figure 24. Example of deviation in read sampling bias across samples.**

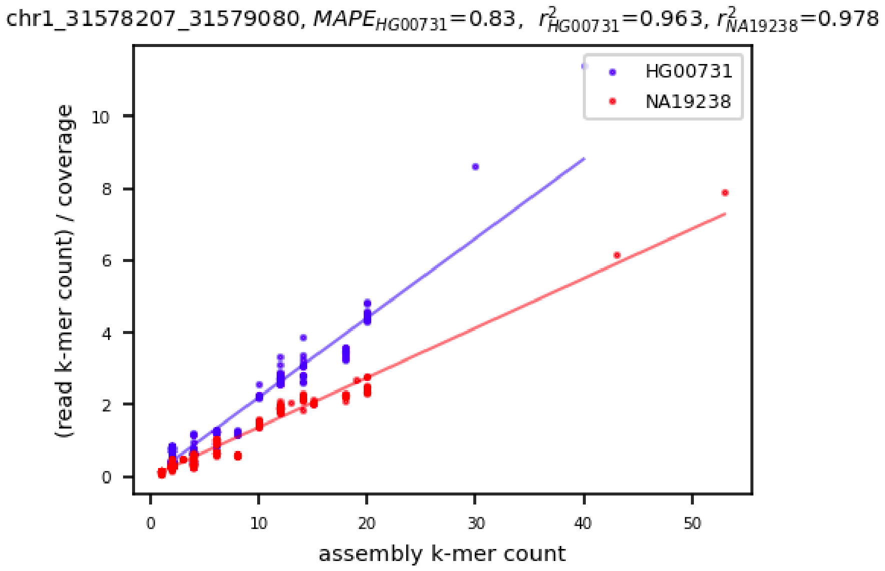


An example locus with high alignment quality but low concordance in read sampling bias between samples. NA19238 has the most similar read sampling bias to HG00731 based on the estimation from 397 control regions and is used to estimate the read sampling bias of this VNTR locus in HG00731. Length prediction error is measured with mean absolute percentage error (MAPE).

**Supplementary Figure 25. Example of under-alignment of orthologous VNTR sequences by pggb.**

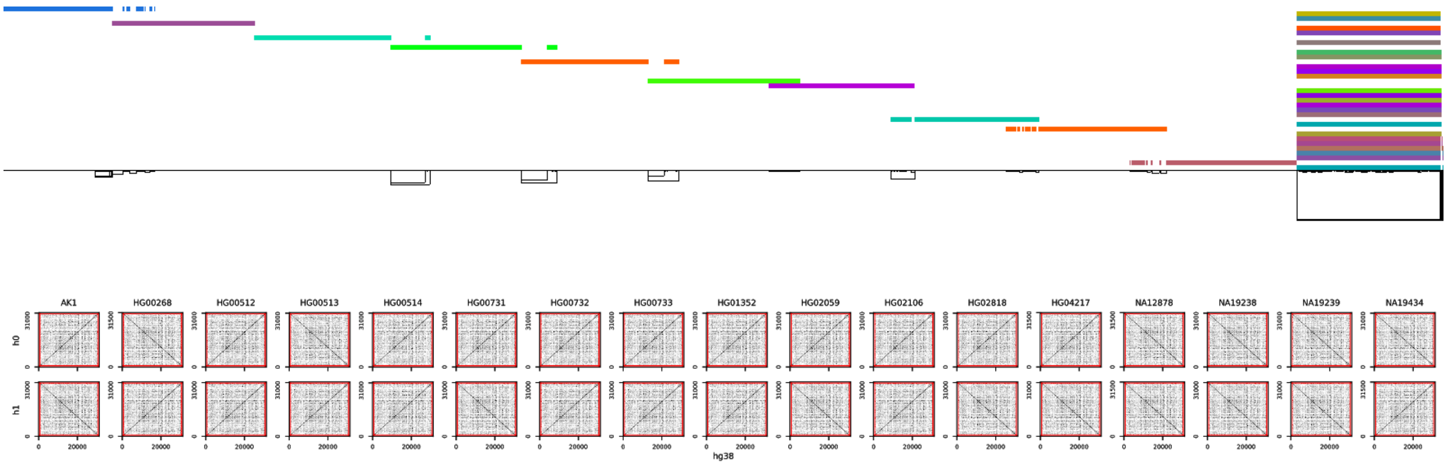

(Top) The multiple sequence alignment result of pggb for 34 VNTR haplotypes at chr12:37898555-37928455 plus 700 bp flanking sequences on each side. (Bottom) The dot plots of all haplotypes against GRCh38.

**Supplementary Figure 26. Misalignment of simulated VNTR reads by bwa.**

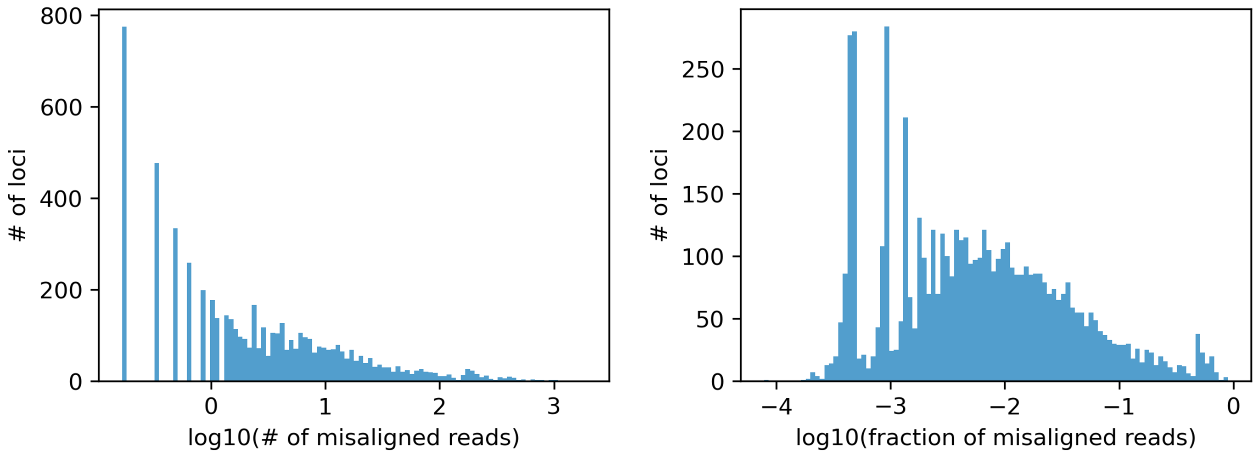


(Left) Number of misaligned VNTR reads averaged across samples. (Right) Fraction of misaligned reads averaged across samples. 32,138 VNTR loci over six genomes, including HG00512, HG00513, HG00731, HG00732, NA19238 and NA19239 were included in this experiment. Loci without misalignments are not shown for clarity. 30x error-free paired-end reads were simulated from the six genomes and each mapped to GRCh38+ALT+decoy+HLA (the hs38DH in bwa) using bwa-mem2 to follow the alignment procedures in the 1KGP and the GTEx project. We define that a read is misaligned if its location is beyond 1 kbp to the boundary of its original VNTR locus.

**Supplementary Figure 27. Misalignment of VNTR reads to GRCh38 rescued by danbing-tk.**

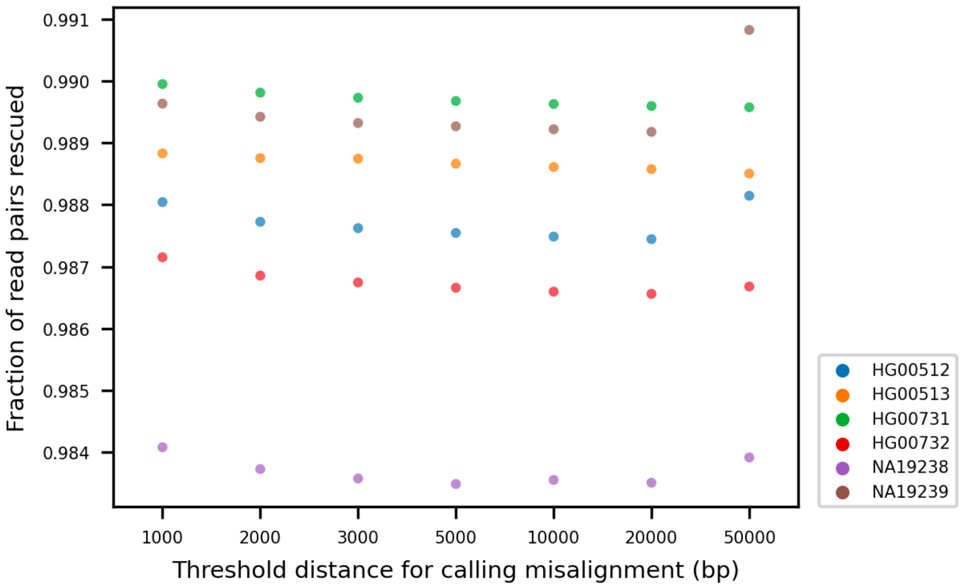


Read pairs misaligned by bwa were extracted and aligned to RPGGs using danbing-tk. A misalignment is called if the distance of any end of the read pair to its original VNTR locus is greater than the threshold. Options "-thcth 50 -cth 45 -rth 0.5" were used for danbing-tk align, same as the setting for genotyping the 1000 and the GTEx genomes.

**Supplementary Figure 28. Relationship between GC content and length prediction error.**

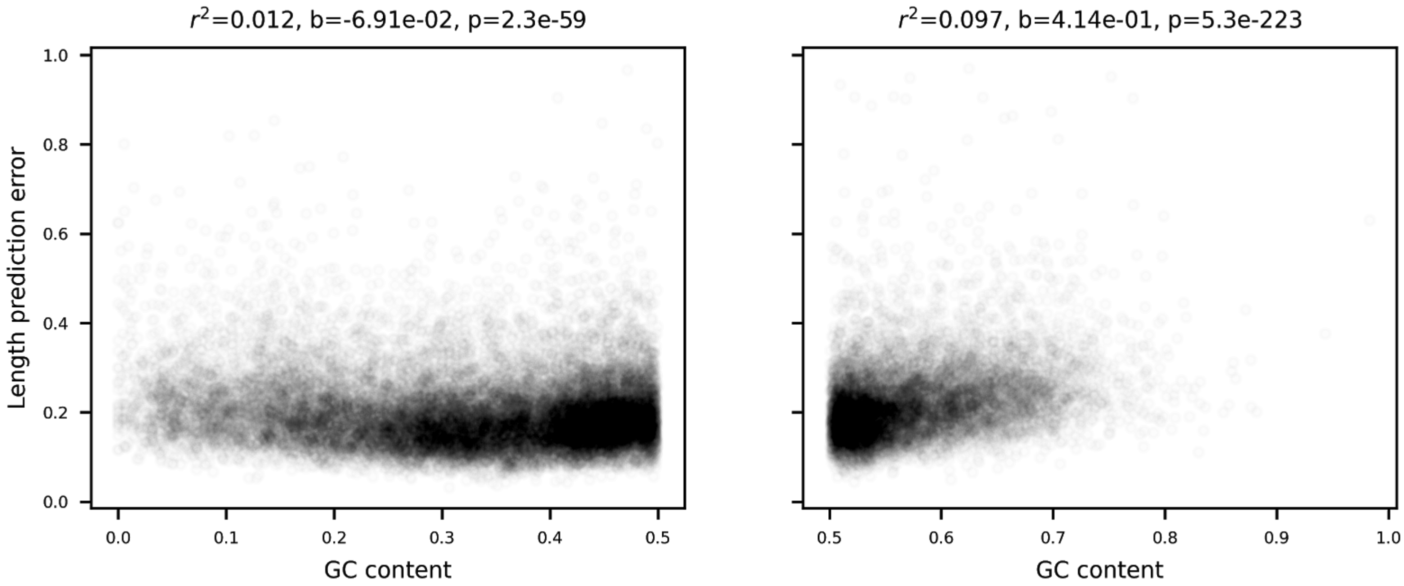


GC contents of the 32,138 VNTRs were measured on GRCh38 using bedtools nuc. Length prediction errors were measured using mean absolute percentage error in the leave-one-out analysis. The r squared, effect size and p value for GC<0.5 (left) and GC>0.5 (right) are shown in the titles.

**Supplementary Figure 29. Relationship between VNTR length and prediction error.**

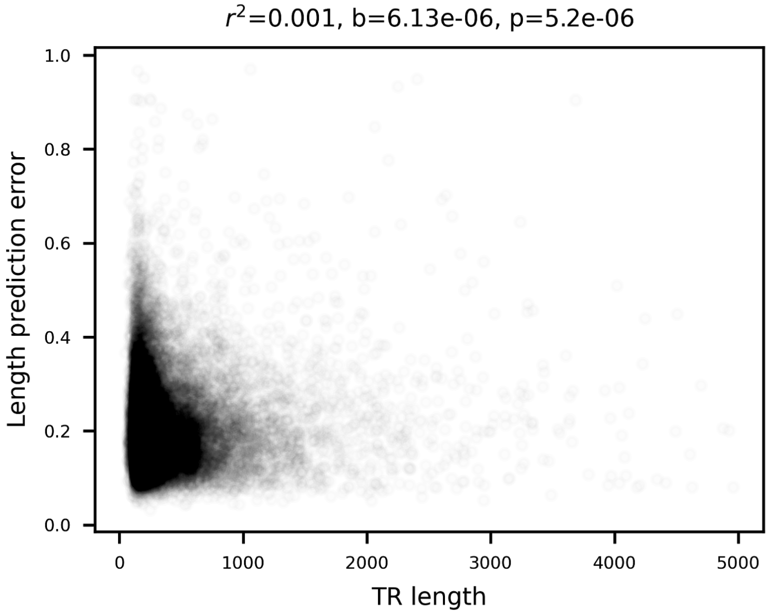


VNTR lengths of 32,138 loci were averaged across 19 genomes. Length prediction errors were measured using mean absolute percentage error in the leave-one-out analysis. The r squared, effect size and p value are shown in the title.

**Supplementary Figure 30. Relationship between eVNTR P-value and prediction error.**

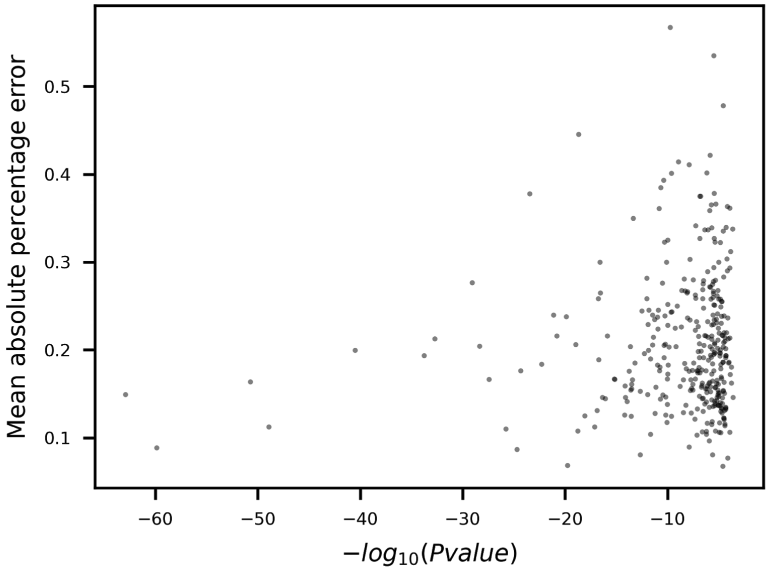


P-values were Bonferroni-corrected. Length prediction errors were measured using mean absolute percentage error in the leave-one-out analysis.

**Supplementary Methods**

Properties of $V_{ST}$

The dosage of a VNTR for an individual $j$ is $x_j \geq 0$. The dosage of an individual $j$ is $x_j$. Consider $P$ populations, with $n_i$ individuals each, with population mean and variance $\mu_i$, and $\sigma_i^2$, and a global mean and variance $\mu_T$ and $\sigma_T^2$ for all individuals.

Population stratification is calculated as $V_{ST} = \frac{\sigma_T^2 - \frac{\sum \sigma_i^2 n_i}{\sum n_i}}{\sigma_T^2}$.

The mean across populations, $u_T$ is calculated as$(\sum \mu_i n_i)/\sum n_i$. The variance is $E(x_j - \mu_T)^2$ for all individuals, and this may be separated out by population as $\sum_i E(x_k^i - \mu_T)^2 n_i / \sum_i n_i$, using $x_k^i$ to denote the $k$th individual in population $i$. The value $E(x_k^i - \mu_T)^2$ may be computed as:

$$\begin{aligned}
E(x_k^i - \mu_T)^2 &= E((x_k^i - \mu_i) + (\mu_i - \mu_T))^2 \\
&= E((x_k^i - \mu_i)^2 + 2(x_k^i - \mu_i)(\mu_i - \mu_T) + (\mu_i - \mu_T)^2) \\
&\text{since } \; E(x_k^i) = \mu_i, E(x_k^i - \mu_i) = E(x_k^i) - E(\mu_i) = 0, \\
&= \sigma_i^2 + E(\mu_i - \mu_T)^2 \\
&= \sigma_i^2 + (\mu_i - \mu_T)^2
\end{aligned}$$

The total population variance $\sigma_T^2$ relative to the population mean, variance, and sizes, and global mean is:
$\sigma_T^2 = \sum_i \sigma_i^2 + (\mu_i - \mu_T)^2)/\sum n_i$.
Replacing this in the calculation of $V_{ST}$ gives:

$$V_{ST} = \frac{1}{\sigma_T^2}(\frac{\sum_i \sigma_i^2 + (\mu_i - \mu_T)^2)}{\sum n_i} - \frac{\sum \sigma_i^2 n_i}{\sum n_i}) \qquad = \frac{\sum(\mu_i - \mu_T)^2}{\sigma_T^2}$$

**Supplementary Table 1. List of HGSVC members.**

| First Name | Last Name | Email | Affiliations |
|---|---|---|---|
| Aaron | wenger | | Pacbio |
| Adam | Mattson | | BC Cancer |
| Alexej | Abyzov | | Mayo Clinic |
| Allison | Regier | | Washington University |

| | | | |
|---|---|---|---|
| Alexej | Hastie | | Bionano Genomics |
| Ali | Bashir | | Icahn School of Medicine at Mount Sinai |
| Amy | Carlough | | The Jackson Laboratory for Genomic Medicine |
| alvaro | Martinez Barrio | | 10X Genomics |
| Anna | Basile | | New York Genome |
| Andre | Corvelo | | new York Genome |
| Arvis | Sulovari | | University of Washington |
| Ashley | Sanders | | EMBL |
| Bernardo | Rodriguez martin | | EMBL |
| Bob | Handsaker | | Broad Institute, Harvard Medical School |
| Brad | Nelson | | University of Washington |
| Can | Alkan | | Bilkent University |
| Charles | Lee | | The Jackson Laboratory for Genomic Medicine |
| Chong | Li | | Temple |
| Christopher | Yoon | | Washington University in St. Louis |
| Chunlin | Xiao | | |
| Conner | Nodzak | | University of North Carolina at Charlotte |
| Daniel | Fordham | | Oxford Nanopore |
| Danny | Antaki | | UCSD |
| David | Porubsky | | |
| Eoghan | Harrington | | Oxford Nanopore |
| Evan | Eichler | | University of Washington |
| Ernest | Lam | | Bionano Genomics |
| Ernesto | Lowy Gallego | | EBI |
| Fabio | Navarro | | Yale University |
| Fereydoun | Hormozdiari | | UC Davis |
| Feyza | Yilmaz | | The Jackson Laboratory for Genomic Medicine |
| Gamze | Gursoy | | Yale |
| Giuseppe | Narzisi | | New York Genome |
| Goo | Jun | | Univ. of Texas Health Science Cetner Houston |
| Haley | Abel | | Washington University in St. Louis |
| Han | Cao | | Bionano Genomics |
| Harrison | Brand | | Harvard |
| Ian | Fiddes | | 10x Genomics |
| Ira | Hall | | Yale |
| Jan | Korbel | | EMBL |
| Jana | Ebler | | |

| | | | |
|---|---|---|---|
| Jason | Chin | | Pacific Bioscience |
| Joel | Rozowsky | | Yale |
| Jonas | Korlach | | Pacific Bioscience |
| Jonathan | Sebat | | University of California San Diego |
| Joyce | Lee | | Bionano Genomics |
| Junjie | Chen | | Temple |
| Kai | Ye | | Xi'an Jiaotong University |
| Katy | Munson | | |
| Ken | Chen | | MD Anderson |
| Kun | Xiong | | Yale |
| Laura Carolyn | Smith | | |
| Letu | Qingge | | UNCC |
| Li | Guo | | Xi'Aan Jiaotong University |
| Li | Ding | | Washington University |
| Lisa | Brooks | | NIH/NHGRI |
| Madhusudan | Gujral | | University of California San Diego |
| Maggi | | | UAB School of Medicine - Birmingham, AL |
| Marc Jan | Bonder | | |
| Mark | Gerstein | | Yale |
| Mark | Batzer | | Louisiana State University |
| Mark | Chaisson | | University Southern California |
| Marta | Byrska-Bishop | | New York Genome |
| Matthew | Wyczalkowski | | Washington University in St. Louis |
| Mike | Smith | | EMBL |
| Mike | Zody | | NY Genome |
| Michael | Schnall-Levin | | 10x Genomics |
| Mike | Talkowski | | Harvard Medical School, Broad Institute, Mass. General |
| Miriam | Konkel | | Clemson |
| Nelson | Chuang | | University of Maryland |
| Nina | Habermann | | EMBL |
| Omar | Shanta | | UCSD |
| Oscar | Rodgriguez | | Icahn School of Medicine at Mount Sinai |
| Paul | Flicek | | EMBL-EBI |
| Peter | Audano | | Univeristy of Washington |
| Peter | Ebert | | Max Plank |
| Patrick | Marks | | 10x Genomics |
| Peter | Lansdorp | | University of British Columbia |
| Qihui | Zhu | | The Jackson Laboratory for Genomic Medicine |

| | | | |
|---|---|---|---|
| Rajeeva | Musunuri | | |
| Rebecca | Serra | rebecca.serra@ | |
| Robel | Dagnow | | USC |
| Ryan | Collins | | Harvard Medical School |
| Ryan | Mills | | University of Michigan |
| Sascha | Meiers | | EMBL Heidelberg |
| Scott | Devine | | University of Maryland |
| Serhat | Tetikol | | Seven Bridges |
| Shamoni | Maheshwari | | 10X Genomics |
| Shantao | Li | | Yale |
| Steve | Sherry | | NCBI |
| Susan | Fairley | | EMBL-EBI |
| Sushant | Kumar | | Yale University |
| Tobias | Marschall | | Heinrich Heine University Dusseldorf |
| Timur | Galeev | | Yale |
| Tobias | Rausch | | EMBL |
| Tonia | Brown | | Univeristy of Washington |
| Uday | Shanker Evani | | New York Genome |
| Vincent | Hanlon | | |
| Virginia | Nunez-Mir | | USC |
| Wan-Ping | Lee | | |
| Wayne | Clark | | New York Genome |
| Weichen | Zhou | | University of Michigan |
| Wen-Wei | Liao | | Wash. University |
| William | Harvey | | Univeristy of Washington |
| Wolfram | Hoeps | | EMBL |
| Xian | Fan | | |
| Xinghua Mindy | Shi | | Temple |
| Xiaofei | Yang | | Xi'an Jiaotong University |
| Xuefang | Zhao | | Harvard |
| Yang | Li | | |
| Zechen | Chong | | UAB School of Medicine - Birmingham, AL |
| Zeid | Hamadeh | | |
| Zev | Kronenberg | | University of Washington |
